# Improving causal effect estimation in multi-ancestry multivariable Mendelian randomization with transfer learning

**DOI:** 10.1101/2025.07.11.664423

**Authors:** Yihe Yang, Xiaofeng Zhu

## Abstract

Multivariable Mendelian randomization (MVMR) has been largely limited to individuals of European ancestry, due to the larger sample sizes available in European genome-wide association studies (GWAS). We introduce MRBEE-TL, one of the first multi-ancestry MVMR methods, which combines transfer learning with bias-corrected estimating equations to improve power in underpowered ancestries and to assess cross-ancestry heterogeneity of disease risk factors. In simulations, MRBEE-TL consistently outperformed MR methods that relied solely on ancestry-specific GWAS data. In real data analyses, MRBEE-TL not only identified ancestry-consistent and ancestry-specific causal effects missed by conventional methods, but also improved power in African and East Asian ancestries. MRBEE-TL is available through the R package `MRBEEX` at https://github.com/harryyiheyang/MRBEEX.

## 1 Background

Mendelian randomization (MR) is an epidemiological method that utilizes genetic variants as instrumental variables (IVs) to infer whether an exposure causally influences an outcome (Bowden et al., 2015, 2016; Zhu, 2021; Sanderson et al., 2022). Since the genotypes of individuals are randomly inherited from their parents and generally do not change during their lifetime, genetic variants are considered to be independent of underlying confounders and hence can be used as IVs to eliminate confounding bias. In addition, Mendelian randomization can be performed by using only genome-wide association study (GWAS) summary statistics (Hemani et al., 2018). The growing availability of GWAS summary data through public repositories such as the GWAS Catalog (MacArthur et al., 2017) and the database of genotypes and phenotypes (dbGaP) (Mailman et al., 2007) has further contributed to the rapid development of MR research. Moreover, MR has been generalized to handle multiple exposures (known as multivariable MR (MVMR)) which enables one to estimate the direct causal effect of each exposure on an outcome (Sanderson et al., 2019; Sanderson, 2021). Existing MVMR methods include multivariable inverse-variance weighting (MVIVW) (Bowden et al., 2016), MVMR-lasso (Rees et al., 2019), genome-wide MR analysis under pervasive pleiotropy (GRAPPLE) (Wang et al., 2021), multivariable MR constrained maximum likelihood (MVMRcML) (Lin et al., 2023), spectral-regularized IVW (SRIVW) (Wu et al., 2024), and MR using bias-corrected estimating equation (MRBEE) (Lorincz-Comi et al., 2024), among many others.

Understanding the heterogeneity of disease risk factors across ancestries is important both for improving the generalizability of findings and for uncovering ancestry-specific disease mechanisms. Given that genetic architectures differ across ancestries (Verma et al., 2024), including different allele frequencies and linkage disequilibrium (LD) patterns, MR is typically performed separately within each ancestry, and the resulting causal estimates are compared across ances-tries (Wang et al., 2022), rather than conducting a universal MR analysis using the trans-ancestry GWAS data. However, this ancestry-stratified strategy has limitations, such as reducing the statistical power for non-European ancestries and increasing the risk of selecting weak instruments. In addition, Hemani et al. (2025) recently demonstrated that MR is more likely to yield inconsistent causal estimates across ancestries if not performed appropriately. Such bias can arise when the MR analysis does not adequately address the LD and allele frequency differences across ancestries, valid instrument selection (Hemani et al., 2025), and potential gene-environment and gene-gene interactions (GEIs and GGIs) (Spiller et al., 2022; Zhu et al., 2024). Thus, it remains challenging to determine whether observed discrepancies in an MR analysis genuinely reflect ancestry-specific disease mechanisms. As a result, there is a growing need for statistical methods that not only account for the aforementioned sources of bias, but also enable joint analysis across diverse ancestries, providing a unified framework to disentangle ancestry-consistent from ancestry-specific disease mechanisms.

Information obtained in European ancestry (EUR) can be borrowed for studies in non-European ancestry, as demonstrated in constructing polygenic risk scores (Ruan et al., 2022) and fine-mapping of causal variants (Yuan et al., 2024). This strategy has recently been introduced into MR as well, aiming to improve the power of causal effect estimation for non-EUR ancestries. For example, Hou et al. (2025) developed trans-ethnic Mendelian randomization (TEMR), one of the first methods designed to enhance causal effect estimation in underrepresented ancestries. However, TEMR is a univariable MR (UVMR) method that relies on the assumption that the estimated causal effects across ancestries follow a multivariate normal distribution, with the correlation matrix estimated from the Z-scores of the selected IVs. As we mentioned above, such an estimated correlation matrix can be sensitive to the selected IVs because of different LD patterns and allele frequencies across ancestral ancestries. When the sample size of the target ancestry is limited, substantial bias may be expected in the estimation of cross-ancestry genetic correlation. IV selection is another significant challenge in MR and is considered to be one of the main sources of spurious, inconsistent causal effect estimates across ancestries. To address this issue, Hemani et al. (2025) systematically compared multiple IV selection strategies and found that fixed effects meta-analysis (FEMA) is the most effective strategy for mitigating cross-ancestry heterogeneity, as it selects regionally optimal IVs based on combined evidence across ancestries. In addition, as demonstrated by Zhu et al. (2024), the genetic effect sizes are much more consistent across ancestries when the genetic variants interacting with environmental factors are excluded. Hemani et al. (2025) recently proposed an efficient MR-based method to detect and remove IVs likely involved in GEI and GGI, thereby enhancing the robustness and generalizability of multi-ancestry MR studies. These improvements in IV selection have been implemented in the CAMeRa platform (Hemani et al., 2025); however, a key limitation is that CAMeRa remains primarily built upon existing UVMR methods.

While improperly conducted MR analyses may yield spurious heterogeneity in causal effects across ancestries, genuine heterogeneity can also arise from environmental and cultural differences, such as diet, climate, and lifestyle (Peterson et al., 2019). For instance, Smith et al. (2024) reported ancestry-specific polygenic mechanisms for type 2 diabetes (T2D), highlighting that East Asians (EAS) exhibit elevated lipodystrophy-related polygenic risk, which contributes to increased T2D susceptibility even at lower body mass index (BMI) levels. Such genuine differences in disease etiology may lead to substantial heterogeneity in causal effect estimates, making their detection an important scientific goal in MR research. For instance, Li and Morrison (2025) conducted a comprehensive investigation using GRAPPLE as the causal effect estimator and identified several exposures with causal effects that differ markedly in magnitude or in direction across ancestries. Accordingly, it is essential to develop MR methods that not only offer adequate statistical power to detect causal effects, but also enable the identification of ancestry-specific heterogeneity in effect sizes.

Here we develop MRBEE-TL, to the best of our knowledge, the first multi-ancestry MVMR method. Specifically, MRBEE-TL extends our previous MVMR approach, MRBEE (Lorincz-Comi et al., 2024), by incorporating transfer learning (Li et al., 2022). MRBEE-TL designates EUR ancestry as the source ancestry and treats East EAS, African (AFR), or other non-EUR ancestries as target ancestries. By leveraging large-scale GWAS data from the source ancestry, MRBEE-TL enhances the statistical power of causal effect estimates in underrepresented populations when effects are consistent across ancestries, while ensuring unbiased estimation when the causal effects are ancestry-specific. Moreover, MRBEE-TL adopts FEMA for IV selection while excluding variants likely involved in GGIs or GEIs to avoid bias.

In our real data analysis of five stroke subtypes, MRBEE-TL not only observed previously reported ancestry-consistent causal effects (Hemani et al., 2025), but also uncovered new biologically plausible causal exposure-outcome pairs of EAS and AFR through transfer learning. In addition, we find that although the dominant causal exposures vary by stroke subtypes, their estimated effect sizes remain consistent across ancestries. These findings suggest that discrepancies in observed causal effects may partly reflect differences in stroke subtype composition across ancestry cohorts, rather than true heterogeneity in causal effect sizes.

## 2 Results

### 2.1 Overview of MRBEE-TL

We introduce MRBEE-TL, to the best of our knowledge, the first multi-ancestry MVMR method that combines MRBEE (Lorincz-Comi et al., 2024) with transfer learning (Li et al., 2022) to improve the causal effect estimation in underpowered ancestry. Generally, transfer learning aims to transfer knowledge from related but distinct datasets to improve the learning performance of a target model. In the context of MR, this involves borrowing information from a well-powered source ancestry to enhance causal effect estimation in a less-powered target ancestry. Here, the target and source MVMR models are formulated as:

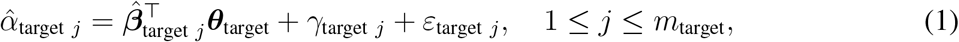

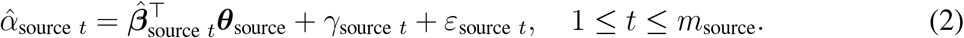

For the target ancestry, 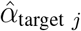 denotes the estimated marginal SNP-outcome association from the outcome GWAS, 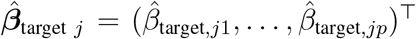 is the vector of SNP-exposure associations from exposure GWASs, ***θ***_target_ is the causal effect vector of interest, *γ*_target *j*_ is the horizontal pleiotropic effect, and *ε*_target *j*_ is the residual error. For thesource ancestry, 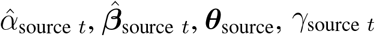, and *ε*_source *t*_ are defined analogously.

An illustration of MRBEE-TL is presented in Figure 1A, and its methodology and implementation are detailed in **Methods** and **Supplementary Materials**. Specifically, MRBEE-TL takes as input a set of GWAS summary statistics for multiple exposures and the outcome in both the source and target populations, along with sample-overlap correlation matrices. Next, MRBEE-TL applies FEMA (Hemani et al., 2025) to jointly select a union set of IVs that are strongly associated with exposures in both the source and target ancestries, using a *χ*^2^-test that accounts for sample overlaps of exposure GWASs via ancestry-specific sample-overlap matrices. In the source population, MRBEE is applied to estimate causal effects, which are then used to get a source causal effect estimate. In the target population, MRBEE-TL is employed to estimate the causal effects by combining several key components: an *𝓁*_2_ loss function, a bias-correction term that accounts for estimation errors in GWAS summary statistics, a penalty function to reduce pleiotropic effects, and a bi-directional selection penalty (BDSP) (Tang et al., 2021) that simultaneously shrinks causal effect estimates toward zero and toward the source causal effect estimates. From a Bayesian perspective, a BDSP can be interpreted as placing a bimodal prior on each target causal effect, centered at zero and the corresponding source estimate (**Methods**). This use of BDSP enables the effective borrowing of information from the source ancestry, thereby enhancing the accuracy of causal effect estimation in the target ancestry. Finally, inference is performed through stability selection to estimate the covariance matrix of target causal effect estimates. The outputs of MRBEE-TL include the causal effect estimates, the corresponding covariance matrix,and the entry-wise posterior inclusion probabilities (PIPs), among many others.

**Figure 1:**
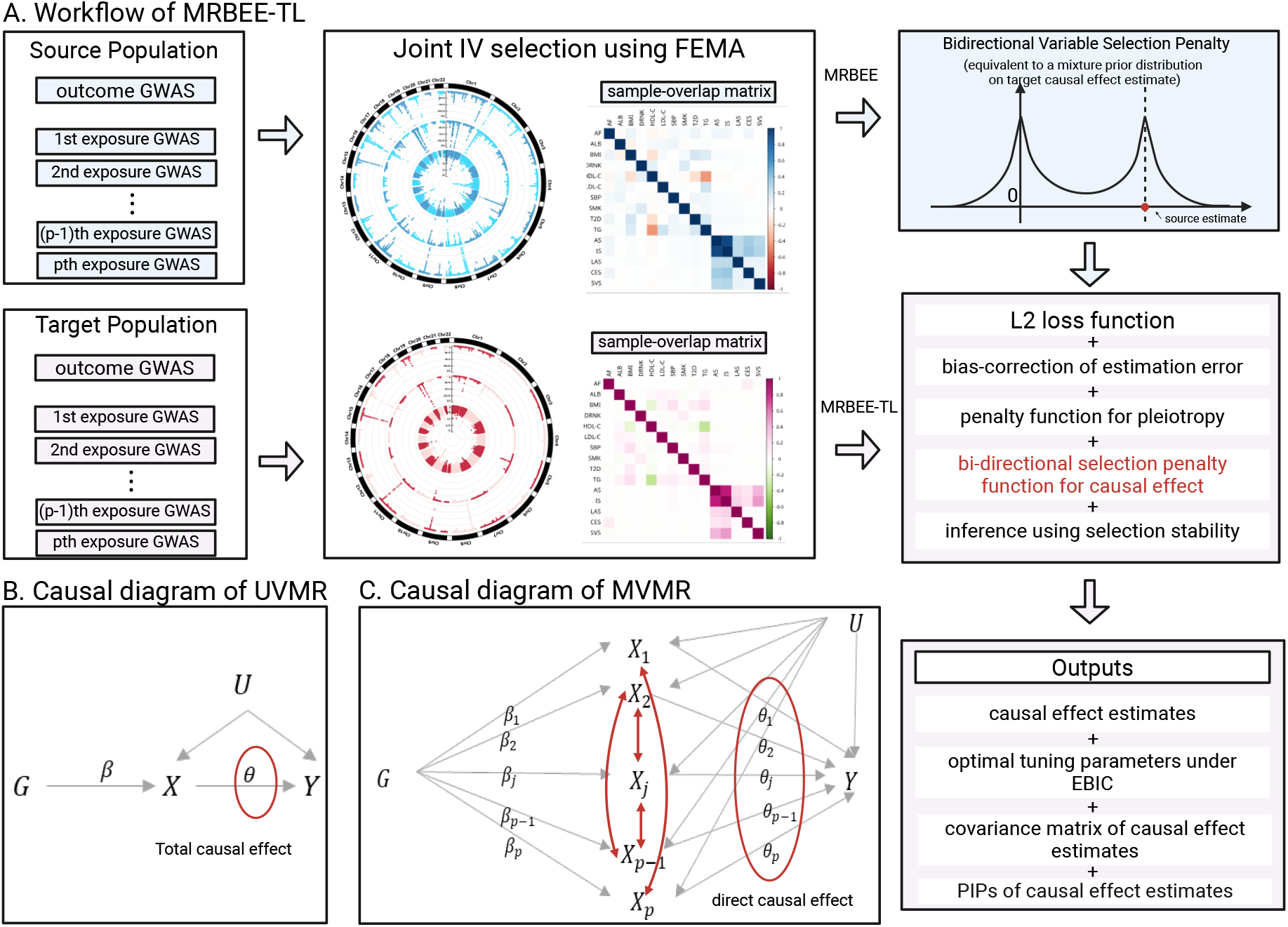
**A.** Illustration of the input components and statistical framework of MRBEE-TL. Each panel represents a key element of the procedure, while arrows indicate the computational flow from inputs to outputs of MRBEE-TL. **B**. Causal diagram of UVMR. **C**. Causal diagram of MVMR. Here, *G* denotes genetic variants, *β* denotes genetic effects from variants to exposures *X, Y* is the outcome, and *U* is the latent confounder. The causal effect of each exposure on the outcome is denoted by *θ*, representing the direct causal effect. All black arrows represent assumed causal relationships in the MR framework. Red arrows highlight the dependency structure among exposures, which MVMR automatically accounts for without requiring explicit modeling.

Compared with traditional MVMR methods restricted to a single ancestry, MRBEE-TL leverages well-powered source summary statistics to construct an empirical prior, thereby enhancing statistical power in underrepresented ancestries. Unlike trans-ethnic UVMR approaches such as TEMR, which estimate total causal effects without accounting for correlations among exposures, MRBEE-TL operates within a multivariable framework and yields direct causal estimates that incorporate the dependency structure among exposures. This distinction is illustrated in Figure 1, where panel B shows the UVMR framework and panel C presents the MVMR structure assumed by MRBEE-TL. Moreover, MRBEE-TL employs a bidirectional separation penalty (BDSP), which allows each exposure to adaptively determine whether to borrow strength from the source ancestry. This avoids imposing unverifiable assumptions about the correlation of causal effects across ancestries, such as equating it to cross-ancestry genetic correlation. Collectively, these features make MRBEE-TL a flexible and powerful tool for disentangling shared and ancestry-specific disease mechanisms.

### 2.2 Simulation

We compared MRBEE-TL against several existing MVMR methods, including the multivariable versions of MR-Median (Bowden et al., 2016), MR-Lasso (Rees et al., 2019), MRcML-BIC (Lin et al., 2023), and MRBEE (Lorincz-Comi et al., 2024). We did not include MRcML-DP in our comparison due to its high computational cost, and we primarily assessed the performance of MRcML-BIC in terms of unbiasedness rather than Type-I error control. In addition, we did not consider non-robust MVMR methods such as IVW (Burgess et al., 2013) and MR-Egger (Bowden et al., 2015), as they are susceptible to bias under horizontal pleiotropy. For traditional MVMR methods, we directly input the GWAS summary statistics from the EAS ancestry. In con-trast, for MRBEE-TL, we first estimated the causal effects in the EUR ancestry using MRBEE, applied hypothesis testing to zero out non-significant entries, and used the resulting vector as the source estimate 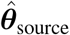 in combination with the EAS summary data. We evaluated method performance across four metrics: (1) root mean square error (RMSE), calculated as 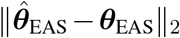; (2) unbiasedness, visualized through boxplots; (3) coverage frequency of the true non-zero causal effect estimates; and (4) Type-I error rate among the null effect estimates. The details of simulation settings are available at **Methods** and **Supplementary Materials**.

As shown in Figure 2, MRBEE-TL consistently produced lower RMSEs than alternative methods, providing an overall summary of its superior performance. This improvement was attributable to two key mechanisms:(1) the incorporation of variable selection to shrink the zero effect estimates to exactly zero, and (2) the use of transfer learning to enforce the ancestry-consistent coordinates of 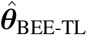 to be exactly the coordinates of 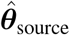. In addition, MRBEE and MRcML-BIC performed similarly, reflecting their shared ability to mitigate estimation error bias through bias correction.

**Figure 2:**
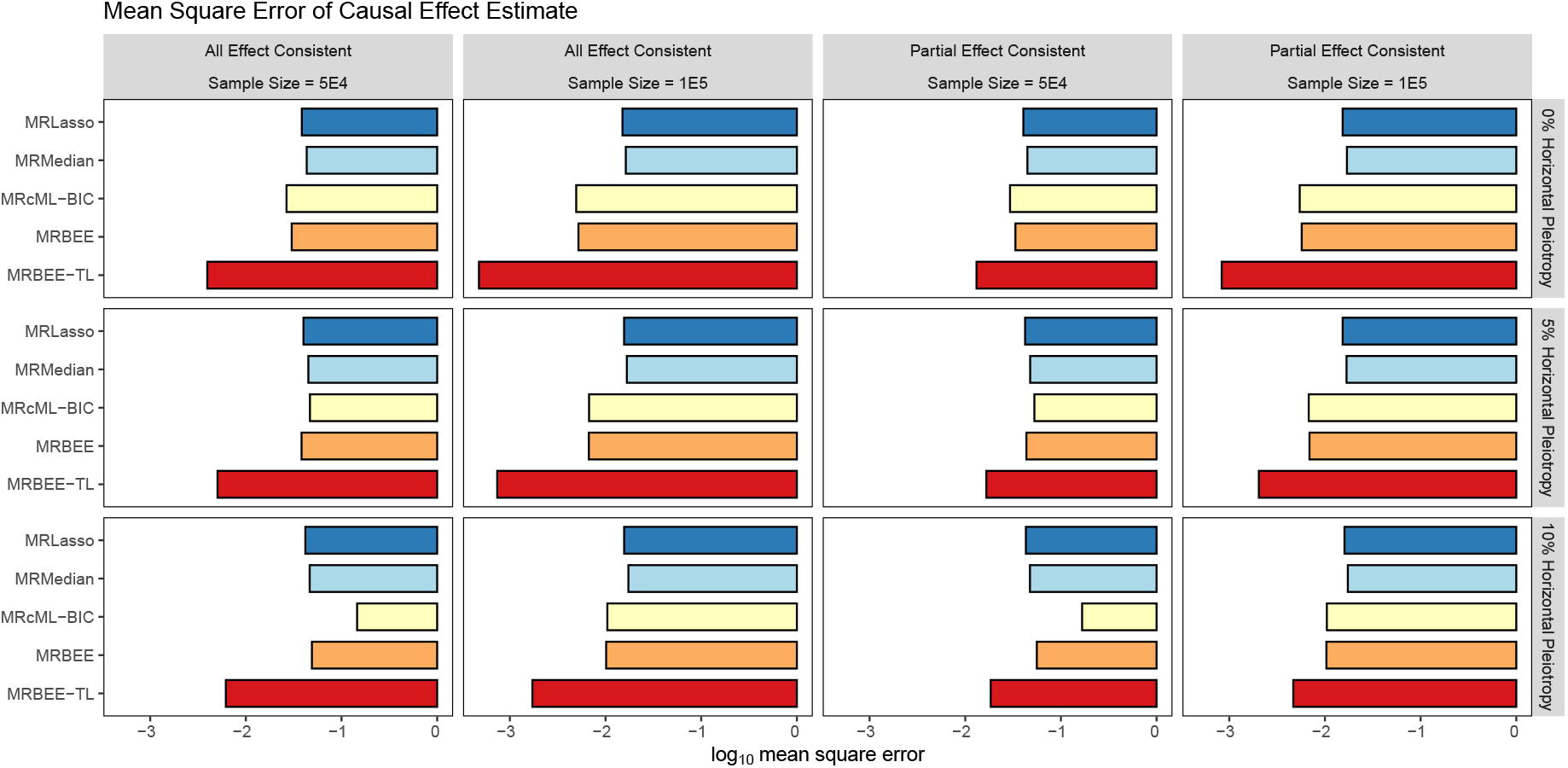
The logarithm of root mean square error (RMSE) of causal effect estimates under full or partial ancestry-consistent effect sizes and different degrees of horizontal pleiotropy. Each panel corresponds to one simulation setting defined by sample size, proportion of horizontal pleiotropy (0%, 5%, or 10%), and whether all or only part of the causal effects are consistent from the source to the target ancestry. Compared methods include MRBEE-TL (proposed), MRBEE, MRcML-BIC, MR-Lasso, and MR-Median. The RMSE is computed as 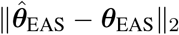. Boxplots summarize the results from 500 simulation repli-cates per setting.

Figure 3 presents boxplots of estimated causal effects for four representative components (*θ*_1_, *θ*_3_, and *θ*_7_) under varying degrees of horizontal pleiotropy and whether or not the causal effects are fully consistent. For *θ*_1_ that had a relatively large effect size and were fully ancestry-consistent, MRBEE-TL achieved substantially lower uncertainty compared to alternative methods. This reflected its ability to correctly recognize these effects as ancestry-consistent and thus borrow information from the source estimate, which was based on a much larger sample size. In contrast, for *θ*_3_ and *θ*_7_, shown in Figure 3C and 3E, MRBEE-TL tended to incorrectly classify some ancestry-specific effects as ancestry-consistent, resulting in over-shrinking toward the EUR estimates and upward bias, when the EAS sample size was small (N = 5E4). This issue became more pronounced with increasing levels of horizontal pleiotropy. Fortunately, this misclassification er-ror dropped sharply when the sample size increased to N = 1E5, suggesting that MRBEE-TL was able to distinguish ancestry-consistent from ancestry-specific effects in large GWAS studies reliably. On the other hand, MRcML-BIC also exhibited visible bias when the proportion of horizontal pleiotropy was large, consistent with patterns previously observed in Lorincz-Comi et al. (2024). Notably, this bias for MRcML-BIC only emerged under high sample-overlap scenarios, a setting particularly relevant for non-European GWAS, where exposures and outcomes are often measured in the same cohort or biobank. As expected, MR-Lasso and MR-Median suffered from severe estimation error bias across all scenarios.

**Figure 3:**
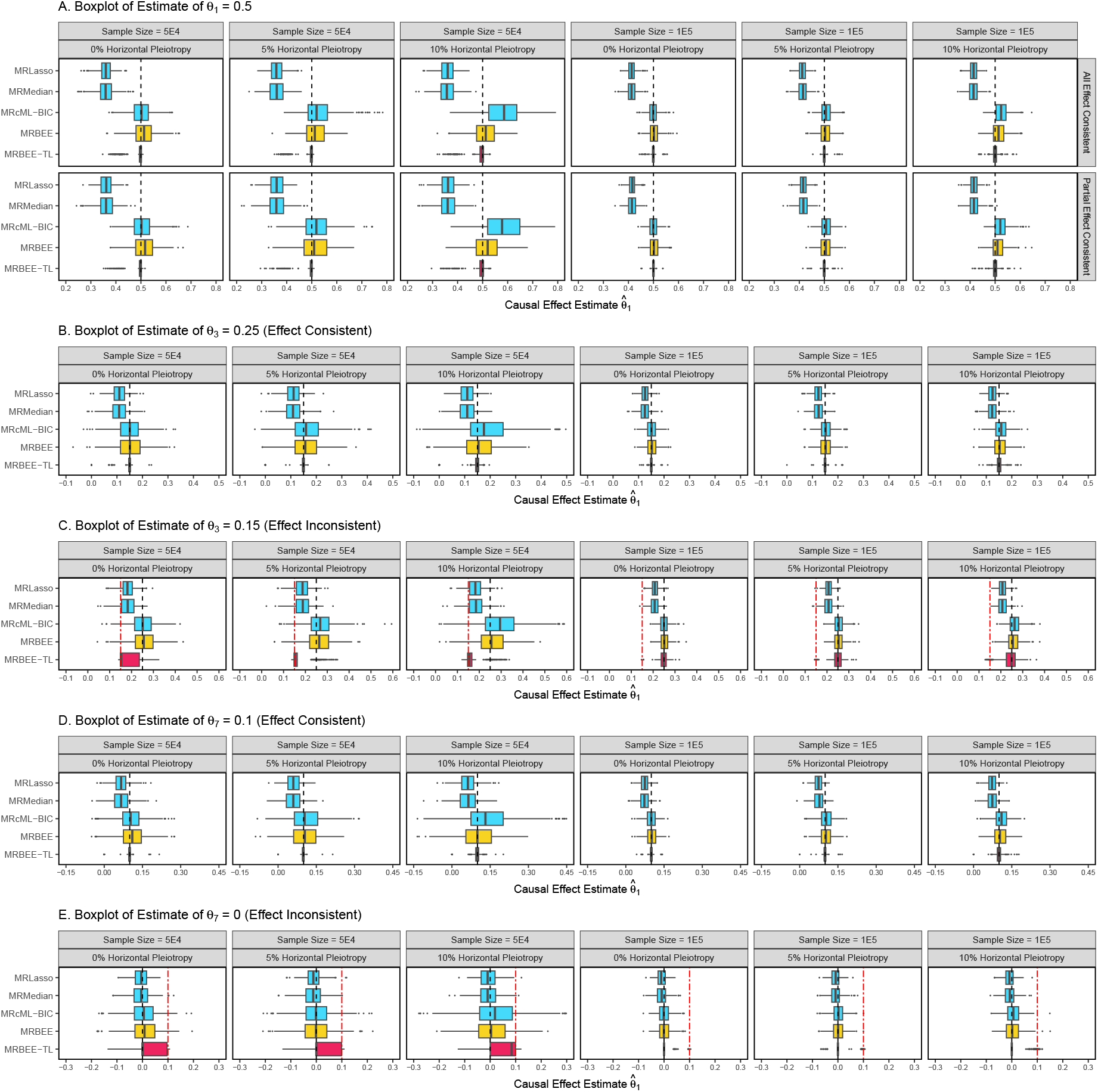
Boxplots of estimated causal effects under full or partial ancestry-consistent effect sizes and different degrees of horizontal pleiotropy. Each panel presents simulation results for one causal component (*θ*_1_, *θ*_3_, or *θ*_7_), under different sample sizes and pleiotropy levels. **A, B** and **D** correspond to causal effects that were fully consistent from EUR to EAS ancestries, whereas **C** and **E** represent inconsistent effects. The black solid lines indicate the true causal effect in the EAS ancestry, and the red dashed lines (in **C** and **E**) denote the corresponding effect estimated from the EUR ancestry, shown only for inconsistent cases. MRBEE-TL, MRBEE, MRcML-BIC, MR-Median, and MR-Lasso were compared. All results were based on 500 replicates per setting.

Figure 4 reports the coverage frequency of 95% confidence intervals for representative causal effects across simulation settings. Coverage frequency refers to the proportion of replicates in which the true effect falls within the 95% confidence interval, and a well-calibrated method should yield coverage near 0.95. MRBEE-TL generally maintained good coverage across most scenarios. However, as the proportion of horizontal pleiotropy increased, coverage declined slightly, most notably in Figure 4C under 10% pleiotropy (last column). When the sample size was small (N = 5E4), MRBEE-TL sometimes misclassified ancestry-specific effects as ancestry-consistent. This led to low coverage in the first three columns of Panel C, where the frequency dropped to around 0.25. MRBEE consistently showed over-coverage across all scenarios, consistent with previous observations (Wu et al., 2024). As expected, MR-Lasso and MR-Median had substantially lower coverage throughout, likely due to uncorrected estimation error bias.

**Figure 4:**
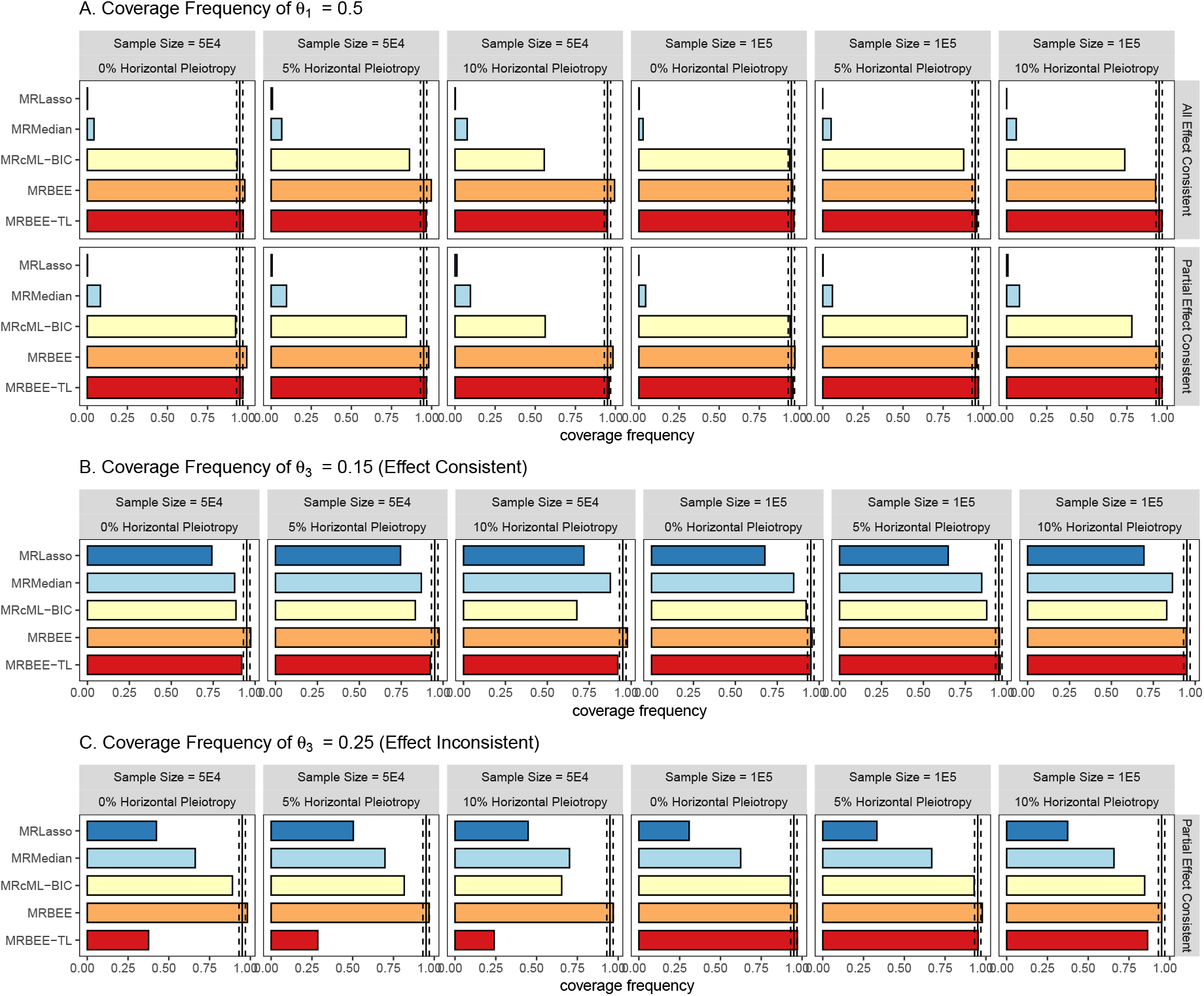
Coverage frequency of 95% confidence intervals for causal effect estimates. Each panel shows the empirical coverage frequency across 500 simulation replicates for selected causal effects (*θ*_1_, *θ*_3_, and *θ*_7_) under varying sample sizes and levels of horizontal pleiotropy. **A** and **B** represent ancestry-consistent effects, while **C** shows a ancestry-specific effect. A well-calibrated method is expected to achieve a coverage frequency close to the nominal level of 0.95. Methods compared include MRBEE-TL, MRBEE, MRcML-BIC, MR-Median, and MR-Lasso. Deviation from the 0.95 line indicates under- or over-coverage due to misclassification of transferability or uncorrected estimation error bias.

Figure 5 evaluates the statistical properties of the compared methods in terms of Type-I er-ror rates and power. Figure 5A and 5C report the empirical rejection frequencies under the null hypotheses *H*_0_ : *θ*_2_ = 0 and *H*_0_ : *θ*_7_ = 0, respectively, while Figure 5B shows the power for detecting a weak but nonzero causal effect (*θ*_7_ = 0.1) and is consistent between EUR and EAS. Overall, MRBEE-TL controlled Type-I errors effectively across most scenarios. Notably, since MRBEE-TL used variable selection to enforce sparsity, it could achieve Type-I error rates well below the nominal 0.05 level in many settings. However, in Figure 5C, where *θ*_7_ was ancestry-specific, MRBEE-TL showed inflated Type-I error rates when the sample size was relativelysmall (N = 5E4), particularly under high levels of horizontal pleiotropy. This reflected the occasional misclassification of ancestry-specific effects as ancestry-consistent in sample-size-limited settings. MRBEE again showed overconservative behavior, with Type-I error rates below 0.05 in many cases, consistent with earlier observations (Wu et al., 2024). MR-Median and MR-Lasso exhibited inflated Type-I errors in nearly all settings, which was likely due to uncorrected estimation error bias, which will raise upward bias when the sample-overlap proportion is large (Lorincz-Comi et al., 2024). Figure 5B further illustrates that when the true causal effect was small but ancestry-consistent, MRBEE-TL achieved substantially higher power than the other methods. This gain was especially visible when N = 5E4, as MRBEE-TL leveraged information from the EUR ancestry to enhance signal detection. MRBEE and MRcML-BIC also achieved moderate power, while MR-Median and MR-Lasso underperformed across all scenarios.

**Figure 5:**
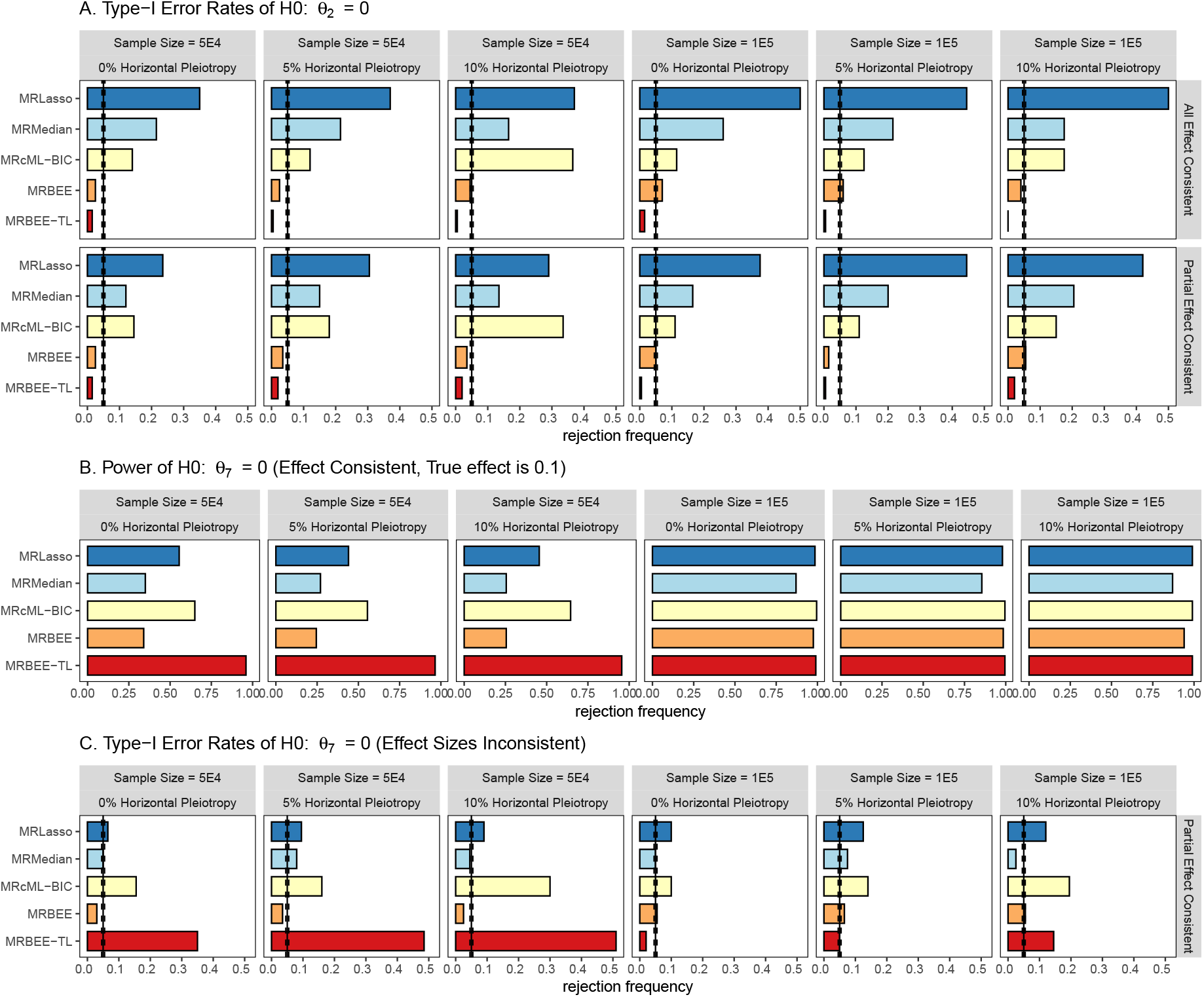
Type-I error rates and power under different simulation settings. Each panel reports the empirical rejection frequency across 500 replicates under varying sample sizes and levels of horizontal pleiotropy.**A** shows Type-I error rates for testing *H*_0_ : *θ*_2_ = 0; **B** presents the power for detecting a weak but ancestry-consistent causal effect *θ*_7_ = 0.1; and **C** evaluates Type-I error rates for testing *H*_0_ : *θ*_7_ = 0, where the effect is population-specific. The horizontal reference line corresponds to the nominal significance levelof 0.05. Compared methods include MRBEE-TL, MRBEE, MRcML-BIC, MR-Median, and MR-Lasso. MRBEE-TL employed variable selection and nonparametric bootstrap inference, allowing for effective control of Type-I errors in most scenarios while also achieving superior power when effects were ancestry-consistent.

### 2.3 Stroke analysis

We applied MRBEE-TL to perform MVMR analyses across five stroke subtypes: all strokes including both ischemic and hemorrhagic strokes (AS), all ischemic stroke (AIS), large-artery atherosclerotic stroke (LAS), cardioembolic stroke (CES), and small vessel stroke (SVS), where the GWAS summary data are provided by Mishra et al. (2022). The exposures included atrial fibrillation (AF), serum albumin (ALB), BMI, drinks per week (DRNK), HDL cholesterol (HDL-C), LDL cholesterol (LDL-C), systolic blood pressure (SBP), smoking initiation (SMK), triglyceride (TG), and T2D, all of which are recognized as common cardiometabolic risk factors across the outcomes of interest. GWAS summary statistics were derived from Graham et al. (2021); Sinnott-Armstrong et al. (2021); Aragam et al. (2022); Saunders et al. (2022); Miyazawa et al. (2023); Chen et al. (2023); Suzuki et al. (2024); Keaton et al. (2024); Verma et al. (2024). The data prepossessing procedure is available at **Supplementary Materials**.

Figure 6 displays the causal effect estimates in East Asian and African ancestries from MRBEE-TL across five stroke subtypes, in comparison to those from MRBEE using ancestry-specific GWAS data. The European causal effect estimates (source) are also present for comparison. Overall, MRBEE-TL resulted in more consistent causal estimates between ancestries than population-specific analysis. For example, MRBEE-TL suggested that ALB was protective and SBP, LDL-C, and SMK are risk factors for AS and AIS in East Asian and African ancestries, which are consistent with European ancestry. In contrast, SMK was not significant in EAS and AFR in ancestry-specific analysis. The causal risk factors varied considerably across the three ischemic stroke subtypes. For LAS, LDL-C and SBP emerged as the primary risk contributors, while ALB and HDL-C were protective factors. Regarding CES, AF was identified as the primary risk factor. T2D was a risk factor in European and East Asian ancestries. The causal factors of CES are less consistent across the three ancestries. For SVS, SBP and TG were consistent risk factors and ALB was protective in the three ancestries. These findings indicate the different causal risk components for stroke subtypes, but the magnitude of causal effects is consistent in the three ancestries for each subtype.

**Figure 6:**
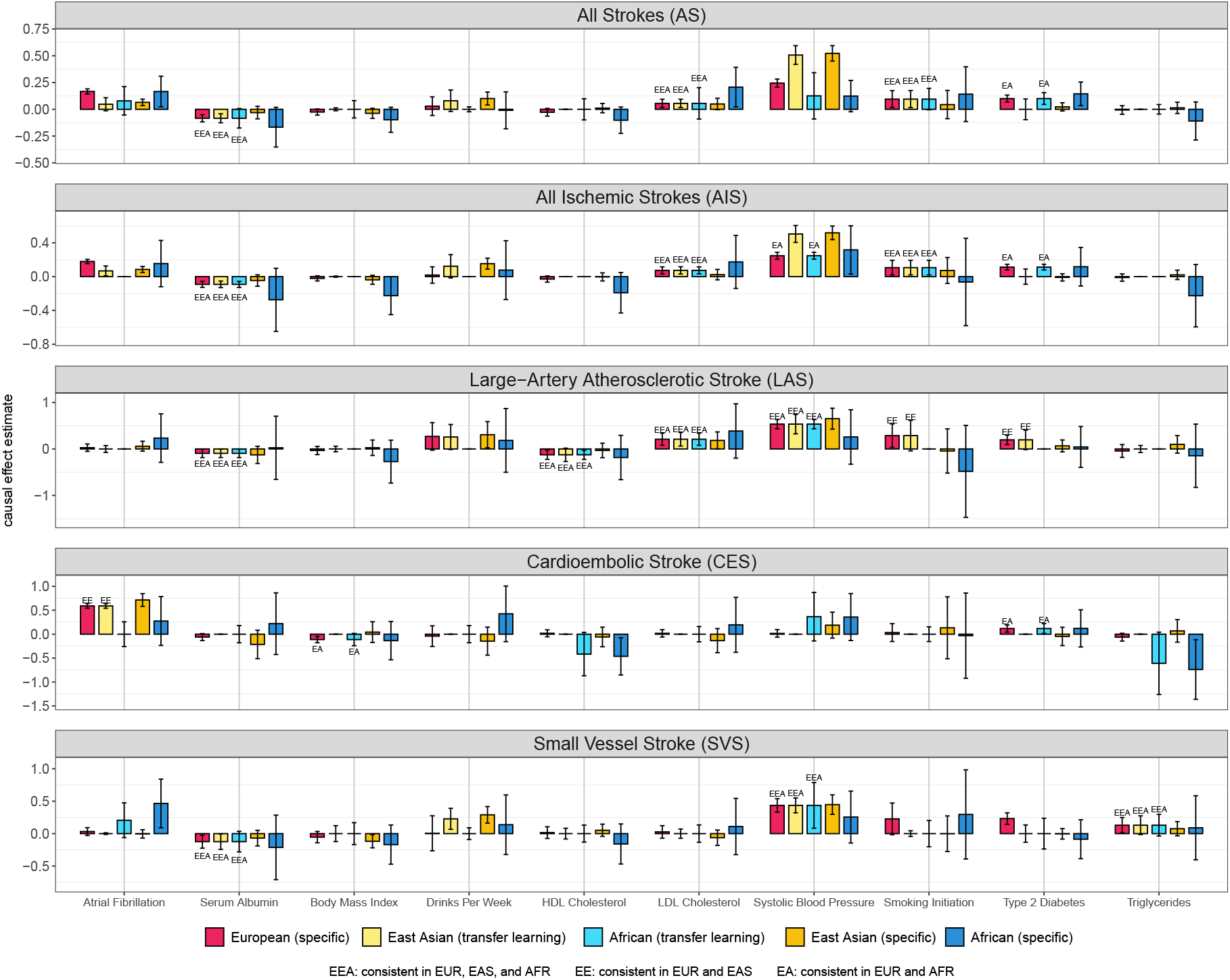
Causal effect estimates of selected exposures on five stroke outcomes across European, East Asian, and African ancestries. Each exposure-outcome pair is evaluated using five settings: European (specific), East Asian (transfer learning), African (transfer learning), East Asian (specific), and African (specific) (by using the original MRBEE). Bar heights represent point estimates of causal effects, with error bars indicating 95% confidence intervals. Annotations “EA”, “EE”, and “EEA” above bars denote consistent causal effect estimates between European and African, European and East Asian, and all three ancestries, respectively, identified by transfer learning.

We observed that multiple causal signals were detected in AFR by MRBEE-TL but were missed by AFR-specific analysis, which can be attributed to the small sample sizes in AFR GWAS. Similarly, in EAS, several causal exposure-outcome pairs, including LDL-C-AS, LDL-C-AIS, SMK-AS, SMK-AIS, HDL-C-LAS, and T2D-LAS, were identified only through MRBEE-TL. These results underscore the value of transfer learning in enhancing statistical power for underrepresented ancestries in GWAS and demonstrate its effectiveness as a scalable extension to traditional MVMR approaches in multi-ancestry analyses. Nevertheless, for CES, BMI was identified as a spurious protective factor in both EUR and AFR ancestries. We currently speculate that this may be due to collinearity among exposures rather than a “novel” exposure with an ancestry-consistent causal effect.

MRBEE-TL was able to identify ancestry-specific risk factors. For example, DRNK was detected as a causal factor for AIS, LAS, and SVS specifically in EAS. In addition, the estimated causal effects of SBP on AS and AIS were much larger in EAS than in EUR or AFR. In contrast, the effect sizes of SBP on LAS and SVS were consistent across ancestries. We interpret this apparent discrepancy as potentially arising from differences in the subtype composition within AS and AIS categories across ancestries, rather than from true heterogeneity in the underlying causal effects, which has been discussed by Hemani et al. (2025).

Figure 7 compares the causal effect estimates obtained from MRBEE-TL and TEMR, where TEMR was performed for each exposure separately. In European ancestry, TEMR identified BMI and TG as significant risk factors and HDL-C as a protective factor for multiple stroke outcomes, while most of these risk-outcome pairs were not significant by MRBEE-TL. The main reason for this discrepancy can be attributed to that MRBEE-TL models all risk factors simultaneously, suggesting that most of the risk factors identified by TEMR do not directly contribute to stroke outcomes. Consequently, TEMR frequently identified BMI and TG as risk factors and HDL-C as protective factors in East Asian and African ancestries. For T2D, TEMR yielded relatively larger effect size estimates than MRBEE-TL, although their 95% CIs always overlapped. In AFR, TEMR identified 7 significant causal exposure-outcome pairs, compared to 12 identified by MRBEE-TL, suggesting that MRBEE-TL is more efficient and powerful than TEMR in utilizing source ancestry information.

**Figure 7:**
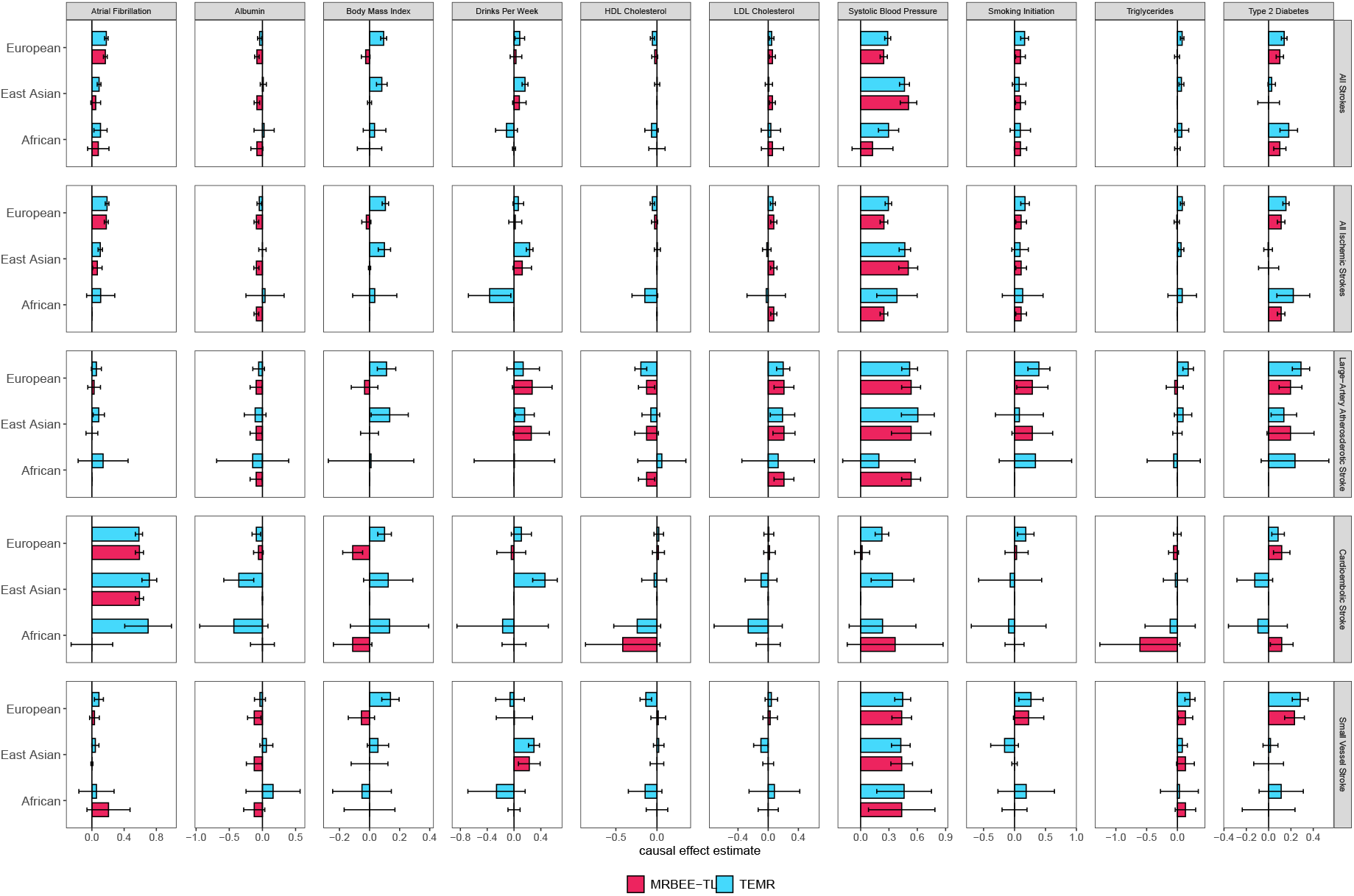
Comparison of causal effect estimates derived from MRBEE-TL and TEMR across three ancestries (for European, the causal effect was derived from MRBEE). Each panel represents one exposure, and within each panel, causal effects on five stroke outcomes are shown. Bars represent point estimates, with error bars indicating 95% confidence intervals. Within each exposure-outcome pair, three ancestry-specific estimates are presented from left to right: European, East Asian, and African American.

## 3 Discussion

Current multi-ancestry MVMR analyses face two major challenges. The first is how to address the limited statistical power caused by the relatively small sample sizes in non-EUR populations. The second is how to distinguish ancestry-consistent causal effects from ancestry-specific ones, which may provide important insights into ancestry-specific disease mechanisms and help inform population-tailored public health policies. To resolve these challenges, we propose MRBEE-TL, an extension of MRBEE that incorporates transfer learning to simultaneously identify ancestry-consistent, ancestry-specific, and non-causal effects. From a mathematical perspective, MRBEE-TL can outperform traditional MVMR approaches as long as a subset of the causal effects is shared between the source and target ancestries (Li et al., 2022), an assumption that is broadly supported by growing empirical evidence (Hemani et al., 2025). Furthermore, MRBEE-TL integrates bias correction and pleiotropy detection, improving robustness in the presence of sample overlap bias and horizontal pleiotropy bias frequently encountered in empirical studies. To reduce bias introduced by inappropriate IV selection, we employ FEMA for IV selection and exclude variants that exhibit GEIs and GGIs.

Our real data analysis suggests that causal factors and their corresponding effect sizes of strokes are generally consistent across ancestral populations. For example, while Hemani et al. (2025) identified LDL-C as causal only for LAS using UVMR analysis, MRBEE-TL indicated its consistent causal effects on AS, AIS, and LAS in all three ancestries, suggesting that LDL-C influences them through shared common causal pathways. Beyond the findings of Hemani et al. (2025), we further identified multiple ancestry-consistent exposure-outcome pairs across the three ancestries, including ALB–AS, ALB–AIS, ALB–LAS, ALB–SVS, SMK–AS, SMK–AIS, HDL-C–LAS, SBP–LAS, SBP–SVS, and TG–SVS. Some effects, such as T2D–AS, T2D–AIS, and T2D–CES, were consistent across EUR and AFR, while others, such as T2D–LAS, SMK–LAS, and CES–AF, were consistent across EUR and EAS. Nevertheless, many of these effects might be undetectable in AFR-specific MR analyses due to the limited sample sizes, highlighting that MRBEE-TL enhances statistical power for underrepresented ancestries.

MRBEE-TL also uncovered significant ancestry-specific heterogeneity. First, as also pointed out by Hemani et al. (2025), some of the observed cross-ancestry discrepancies in causal effect estimates, particularly for composite outcomes like all stroke or all ischemic stroke, may reflect differences in subtype composition rather than true heterogeneity in causal mechanisms. For instance, the estimated effects of SBP on AS and AIS were consistently smaller in EUR compared to EAS. In contrast, SBP exhibited broadly consistent effects across ancestries for LAS and SVS. Indeed, the total number of EUR cases for these specific subtypes was only 24,114, substantially less than the 73,652 AS cases and 62,100 AIS cases, which may partly explain their different contributions to overall stroke risk across ancestries. In addition, our analysis identified DRNK as a potential ancestry-specific risk factor for stroke in EAS, a finding that, to our knowledge, has rarely been discussed in the literature. Further investigation of ancestry differences in the genetic architecture of alcohol consumption and its downstream disease mechanisms may offer important insight and represent a promising direction for future research.

In the comparison between TEMR and MRBEE-TL, we observed a fundamental distinction between the two methods: MRBEE-TL is an MVMR method that estimates the direct causal effect, whereas TEMR is a UVMR method that estimates the total causal effect, which is the sum of the direct causal effect and the medication effects through other exposures. While both quantities are interpretable within a causal inference framework, they reflect different aspects of the underlying biology (Sanderson et al., 2019). In our opinion, an MVMR method could be more suitable than a UVRM method if the goal is disentangling the independent contributions of multiple, potentially correlated exposures. In addition, in this comparison, MRBEE-TL identified more causal exposure–outcome pairs for AFR (12 vs. 7), possibly because its use of transfer learning allows for more effective borrowing of information from high-powered source populations.

While MRBEE-TL adopts the FEMA proposed by Hemani et al. (2025) for IV selection, it is noted that this IV selection strategy was originally developed for UVMR analysis. MRBEE-TL extends it to MVMR by jointly testing the association of each variant with multiple exposures across ancestries with a *χ*^2^-statistic. For example, in our analysis involving three populations and ten exposures, the joint test was conducted using a 30-degree-of-freedom *χ*^2^-distribution. Furthermore, MRBEE-TL excludes IVs with potential GEIs through an MR-GxE framework, which is particularly important given the differing environment distributions across ancestries (Zhu et al., 2024). Finally, MRBEE-TL builds upon the MVMR framework, representing a significant advance over TEMR and the platform CAMeRa, both of which are based on the UVMR framework.

MRBEE-TL also has several limitations. First, MRBEE-TL treats the source causal effect estimates as fixed inputs, without jointly updating them alongside the target estimates. As a result, for exposures with truly ancestry-consistent effects, MRBEE-TL does not leverage information from the target ancestry, leading to a potential loss in statistical power. A unified transfer learning framework that jointly estimates causal effects across multiple ancestries could address this limitation. Second, although MRBEE-TL might have selected several additional exposure-outcome pairs via a SuSiE-based variable selection procedure, such as the HDL-C–AS and SMK–AS pair for AFR, their 95% confidence intervals were wide and included zero, suggesting a lack of statistical significance. Further investigation with larger sample sizes is necessary to make a clear causal conclusion. In addition, for certain exposure-outcome pairs, such as SBP–SVS in AFR, the confidence intervals were noticeably wider than in EUR, despite showing consistent effect sizes. Both phenomena likely stem from the same source: in some replicates of stability selection, the exposure was not consistently classified as ancestry-consistent. Furthermore, as Hemani et al. (2025) emphasized, IV selection and other improper quality controls in multi-ancestry MR are subject to several inherent challenges. MRBEE-TL, like other MR methods, relies on the validity of IV selection and cannot inherently correct for biases arising from suboptimal IV selection. Therefore, the ancestry-specific heterogeneity identified by MRBEE-TL should also be interpreted with caution, as it may partially reflect biases introduced during IV selection rather than true biological differences.

## 4 Conclusion

MRBEE-TL extends MRBEE with transfer learning to estimate direct, multivariable causal estimates, thereby augmenting power in ancestries with limited GWAS data while objectively partitioning shared versus ancestry-specific effects. Across extensive simulations and a real data analysis regarding strokes, MRBEE-TL consistently surpassed leading MVMR and trans-ethnic univariable MR methods, revealing additional, biologically credible risk factors in AFR and EAS ancestries. In summary, by enabling rigorous, scalable causal inference across diverse populations, the open-source MRBEE-TL platform advances genomic medicine and promotes equity in genetic epidemiology.

## 5 Methods

### 5.1 Multivariable Mendelian randomization model

Let ***g***_*i*_ = (*g*_*i*1_, …, *g*_*im*_)^⊤^ be an (*m ×* 1) genotype value vector of *m* genetic variants, ***x***_*i*_ = (*x*_*i*1_, …, *x*_*ip*_)^*⊤*^ be an (*p ×* 1) vector representing *p* exposures, and *y*_*i*_ be an outcome. Here, *m* is the number of IVs, which is usually the number of independent loci with *p*-values reaching the genome-wide significant level. The MVMR model is below:

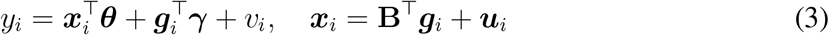

where **B** = (***β***_1_, …, ***β***_*m*_)^*⊤*^ is an (*m × p*) matrix of genetic effects on exposures with ***β***_*j*_ = (*β*_*j*1_, …, *β*_*jp*_)^*⊤*^ being an (*p×* 1) vector, ***θ*** = (*θ*_1_, …, *θ*_*p*_)^*⊤*^ is an (*p×* 1) vector of causal effects of the *p* exposures on the outcome, ***γ*** = (*γ*_1_, …, *γ*_*m*_)^*⊤*^ is an (*m×* 1) vector, known as the horizontal pleiotropy, and ***u***_*i*_ and *v*_*i*_ are the noise terms. We assume that the total number of exposures *p* is fixed and the causal effect ***θ*** is fixed and bounded. The genetic variant *g*_*ij*_ is standardized so that E(*g*_*ij*_) = 0 and var(*g*_*ij*_) = 1, and all IVs are in linkage equilibrium (LE), i.e., cov(*g*_*ij*_, *g*_*ik*_) = 0 for *j/*= *k*. Substituting for ***x***_*i*_ in (3), we obtain the reduced form: *y*_*i*_ = ***α***^*⊤*^***g***_*i*_ + ***θ***^*⊤*^***u***_*i*_ + *v*_*i*_, where ***α***_*i*_ = (*α*_1_, …, *α*_*m*_)^*⊤*^, and the genetic effects of exposures and outcome have the following relationship:

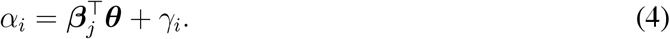

In practice, neither the outcome genetic effects *α*_*j*_ nor the exposure-specific genetic effects ***β***_*j*_ are directly observed; instead, they are usually estimated from GWAS. 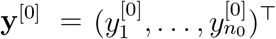 be the sample vector from outcome GWAS, 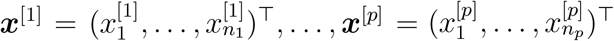 be the sample vectors of the 1st, …, *p*th exposure GWAS cohorts, and 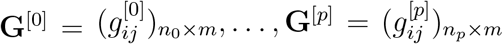 be the sample matrices of *m* genetic variants of the outcomeand 1st, …, *p*th exposure GWAS cohorts. The sample size of the *s*th GWAS cohort is *n*_*s*_, and the overlapping sample size between the *s*th and the *k*th GWAS cohorts is *n*_*sk*_. The estimates of 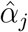 and 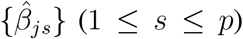 are thus yielded by 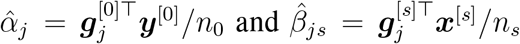,where 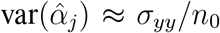 and 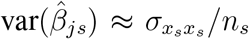. Then the GWAS summary data are formed by 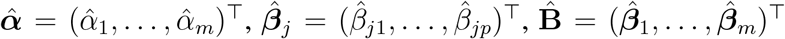, the related standard errors (SEs), the *p*-values, and SNPs information. The summary-statistics-based MR model refers to:

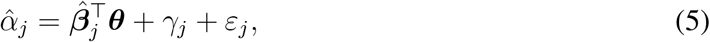

where 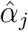 and 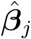 are estimated from outcome and exposure GWAS for *j*th IV, and *ε*_*j*_ represents the residual of this regression model. In addition, the GWAS summary data are usually standardized, i.e., 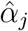 and 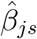 by 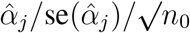 and 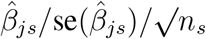, to remove the minor allele frequency effect (Lorincz-Comi et al., 2024).

### 5.2 Mendelian randomization using transfer learning

Recall that MRBEE-TL considers the following target and source model:

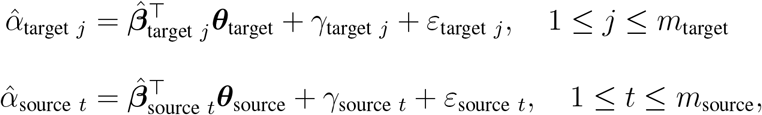

which has been demonstrated in (1)-(2). MRBEE-TL does not assume that the source and target models share the same set of causal instruments. Therefore, we use different index variables *j* and *t* and allow the number of instruments *m*_target_ and *m*_source_ to differ across populations. In addition, for notational simplicity, we omit the subscript “target” in 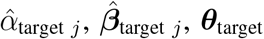, and *m*_target_ in the remainder of the paper.

MRBEE-TL aims to estimate ***θ*** more precisely by borrowing information from a source causal effect estimate 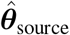, which is typically obtained from a European ancestry. Specifically, MRBEE-TL considers the following penalized minimization:

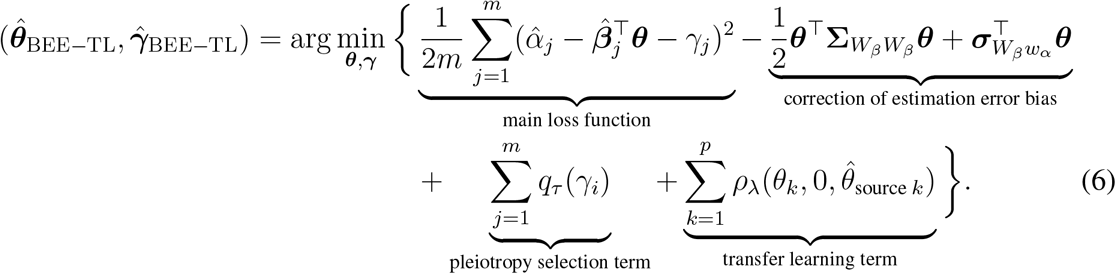

In this minimization, there are three terms in addition to the main loss function. The first term is 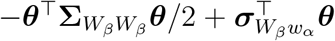, which is just the correction term of estimation error bias used in MRBEE (Lorincz-Comi et al., 2024). The second penalty is ∑_*j*_ *q*_*τ*_ (*γ*_*i*_) with a tuning parameter *τ*, which is employed to identify the horizontal pleiotropy that resemble outliers (Zhao et al., 2020). In practice, lasso is one of the most well-known penalties (Tibshirani, 1996), and MR-Lasso adopts it to identify invalid IVs with large horizontally pleiotropic effects (Rees et al., 2019). Whereas, other variable selection penalties such as the minimax concave penalty (MCP) (Zhang, 2010) can also be used here.

The third penalty *ρ*_*λ*_(*θ, ϑ*_1_, *ϑ*_2_) is the key component that enables MRBEE-TL to distinguish ancestry-consistent and ancestry-specific causal effects. This penalty, known as the bidirectional separation penalty (BDSP) (Tang et al., 2021), encourages each effect to be either zero or similar to its source estimate, whichever makes more sense given the data. Formally, it is defined as:

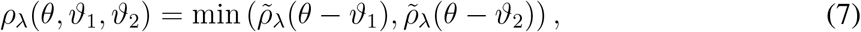

where 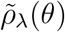 is a common variable selection penalty such as lasso, and *ϑ*_1_ and *ϑ*_2_ represent twodirections of penalization. For the *k*th target causal effect *θ*_*k*_, we set *ϑ*_1_ = 0 to shrink small or non-causal effects toward zero, and set 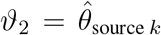 to pull ancestry-consistent effects toward the source estimate. By using BDSP, MRBEE-TL can achieve the following two goals:

1. **Identifying ancestry-consistent effects:** When a target causal effect is consistent with the source causal effect, i.e., 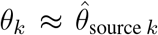, BDSP encourages alignment with 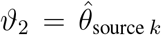, improving estimation of underpower ancestry via transfer learning.
2. **Detecting ancestry-specific effects:** When a target causal effect *θ*_*k*_ differs from the source causal effect 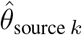, BDSP avoids forcing similarity, allowing data-driven identification of ancestry-specific heterogeneity.

Note that we define exposure as ancestry-consistent in a target ancestry if its true causal effect size is identical to that in the source ancestry. In practice, when the estimated causal effect in the target ancestry is statistically consistent with the source estimate, we say that we have identified an ancestry-consistent effect.

From the Bayesian point of view, the penalty function can be interpreted as the negative log density of a prior distribution on the parameter. Specifically, we consider *p*_***λ***_(***θ***) *∝ −* log *f*_***λ***_(***θ***), where *f*_***λ***_(***θ***) denotes the prior density on the causal effect vector ***θ***_*j*_. When a standard Lasso penalty is used, the corresponding prior is a Laplace distribution: *f*_*λ*_(*θ*) *∝* exp(*−λ*|*θ*|), which encourages sparsity by placing most of the prior mass around zero (Park and Casella, 2008). In our bi-directional selection setting, we instead define a prior density proportional to

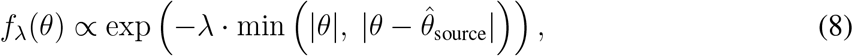

which is centered at both 0 and the source ancestry estimate. The pattern of this prior distribution can be found in Figure 1. To tune the shrinkage parameter *λ*, we use a data-driven criterion such as the extended Bayesian information criterion (EBIC) (Chen and Chen, 2008), which balances model fit and complexity while accounting for the large model space, allowing the method to select more informative, weak-to-moderate signals that would otherwise be overshrunk under fixed penalization. This adaptive selection improves statistical power while maintaining interpretability. Similar to the Lasso, other commonly used variable selection penalties also admit corresponding Bayesian interpretations (Moreno et al., 2015).

### 5.3 Instrument Selection

We adopt FEMA (Hemani et al., 2025) to select the IVs. Let *K* be the total number of ancestries under study; in our stroke application, we have *K* = 3: one source (EUR) and two targets (EAS and AFR). For each SNP *j* and exposure block of dimension *p*, we stack the marginal effect estimates from the *k*-th ancestry in 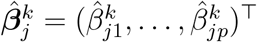 and compute the test statistic

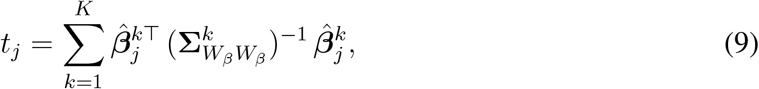

where 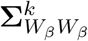 is the estimation error covariance matrix of 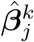. Under the joint null hypothesis 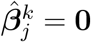 for all *k* = 1, …, *K*, the statistic in (9) follows a chi-squared distribution with

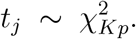

FEMA selects the *j*th variant as IV if *t*_*j*_ exceeds the genome-wide significance threshold. In addition, the estimation procedure of 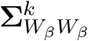 can be found in Lin et al. (2023); Lorincz-Comi et al. (2024)

We excluded the IVs with potential GEI using two procedures. We removed IVs located near the GxE interaction loci (e.g., within 500 KB of the leading variants), based on published studies such as Sung et al. (2018); Zhu et al. (2024). Second, inspired by Zhu et al. (2024), where their Figure 5 reveals that variants exhibiting GxE interactions tend to have highly inconsistent marginal effects across ancestries, we performed MR analysis to detect pleiotropic outliers by treating European ancestry’s GWAS as the exposure and the other’s GWAS as the outcome (Zhu, 2021; Lorincz-Comi et al., 2024). Variants identified as horizontal pleiotropic outliers in this setting were excluded from the analysis.

### 5.4 Implementation

MRBEE-TL iteratively updates each parameter in the minimization (6) using a profile-likelihood (Murphy and Van der Vaart, 2000). Specifically, let ***θ***^(*t*)^ and ***γ***^(*t*)^ be the estimates of ***θ*** and ***γ*** at the *t*th iteration. Following the procedure of Tang et al. (2021), MRBEE-TL will construct a working parameter 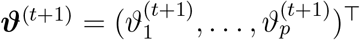 with the *k*th entry being

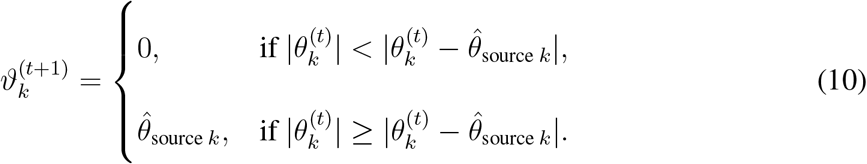

In other words, MRBEE-TL clusters each coordinate of ***θ***^(*t*)^ around two possible centers: 0 and the corresponding entry in the source estimate 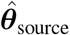. Based on the working center vector ***ϑ***^(*t*)^, we define the difference parameter ***δ*** = ***θ*** *−* ***ϑ***^(*t*)^, which captures deviation from either 0 or ancestry-consistent causal effect, thereby allowing the algorithm to adaptively select plausible causal effect during each iteration given the kth exposure. Accordingly, the updated difference parameter ***δ***^(*t*+1)^ is obtained by solving the following optimization problem:

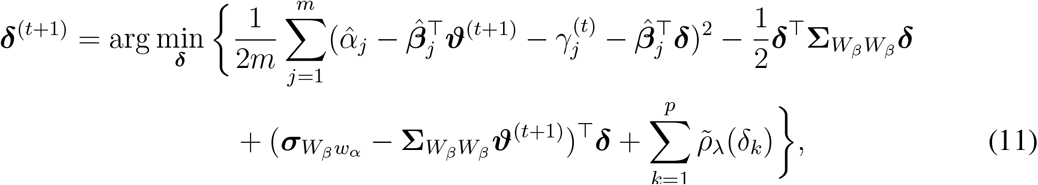

where 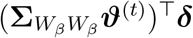 is an additional correction term of estimation error bias due to the inclusion of ***ϑ***^(*t*+1)^. That is, the introduction of the working center ***ϑ***^(*t*)^ transforms the non-common BDSP *ρ*_*λ*_(*θ, ϑ*_1_, *ϑ*_2_) into a common variable selection penalty 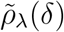, which can be efficiently han-dled by existing algorithms. After obtaining ***δ***^(*t*+1)^, ***θ*** is then updated by ***θ***^(*t*+1)^ = ***δ***^(*t*+1)^ + ***ϑ***^(*t*+1)^. The update of ***γ*** follows standard procedures widely used in the literature; see, e.g., *cis*-MRBEE (Yang et al., 2025). Conditioning on ***θ***^(*t*+1)^, the original minimization in (6) reduces to:

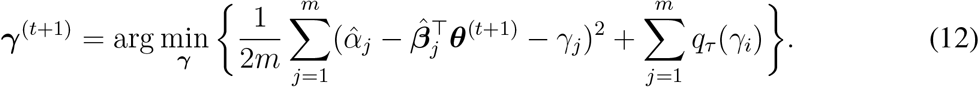

For a broad class of penalty functions *q*_*τ*_ (·), ***γ***^(*t*+1)^ admits a closed-form expression. The detailed procedure for solving the minimization (6) is described in the **Supplementary Materials**.

Although we use lasso as an example to illustrate the principle of BDSP, we actually use SuSiE (Wang et al., 2020; Zou et al., 2022) as the penalty 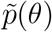 in the minimization (11) to identify non-zero entries of ***δ***. The prior distribution used in SuSiE can be interpreted as the spike-and-slab distribution (Wang et al., 2020), which has been shown to outperform lasso in terms of unbiasedness and model selection consistency (Ročková and George, 2018). In SuSiE, although multiple tuning parameters are available, we only tune the number of single effects *L*, which we refer to as *λ* in 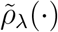, and use the default settings for the remaining parameters. However, because SuSiE typically assumes a small number of single effects *L*, it is not well suited for selecting horizontal pleiotropy. Therefore, we turn to use MCP (Zhang, 2010) as *q*_*τ*_ (*γ*) in the minimization (12). In practice, MRBEE-TL adopts the EBIC (Chen and Chen, 2008) to select both the optimal number of single effects *L* in SuSiE and the tuning parameter *τ* in MCP.

To estimate the covariance matrix of 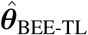, MRBEE-TL uses a selection stability method (Meinshausen and Bühlmann, 2010), where a [0.5*m*] number of the independent IVs are resampled without replacement. According to Meinshausen and Bühlmann (2010), the sample size of [0.5*m*] is chosen as it resembles most closely the bootstrap while allowing computationally efficient implementation, and our previous work (Yang et al., 2024) shows that selection stability works well for estimating the covariance matrix of estimates. Based on the resampled IVs, MRBEE-TL re-estimates the causal effects using the optimal tuning parameters selected by EBIC. The final covariance matrix is computed based on the empirical variation across the replicates. We recommend setting the number of resampling iterations to 1,000, and this choice is used consistently in both the simulation studies and real data analyses. In particular, for estimates deemed consistent with the source ancestry, the final covariance matrix incorporates the covariance of the source estimates; that is,

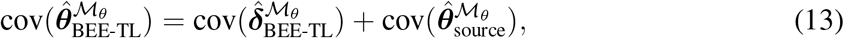

where ℳ_*θ*_ is an index set of exposures whose effect sizes are consistent between source and tar-get population, 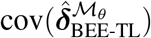 is the bootstrap-estimated covariance matrix of the differences, and 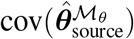 is the covariance matrix of the corresponding source estimate. In other words, the uncertainty of the source estimate is jointly incorporated into MRBEE-TL. In addition, we recommend applying a lenient significance threshold (e.g., P*<*0.05), below which the causal effect estimate in the source ancestry is forcibly set to zero. This helps avoid misleading interpretations of ancestry-consistent effects that may arise from weak or non-significant signals. See the **Supplementary Materials** for implementation details.

### 5.5 Simulation

We conducted simulation studies to evaluate the performance of MRBEE-TL. We generated GWAS summary statistics for *p* = 10 exposures, of which three were designed to have nonzero causal effects in the source (EUR) and target (EAS) ancestries. Besides, we considered three configurations for the true causal effect vectors: (i) fully ancestry-consistent effects, where ***θ***_EAS_ = ***θ***_EUR_ = (0.5, 0, 0.15, 0, 0.1, 0, 0.1, 0, 0, 0)^*⊤*^; (ii) partially ancestry-consistent effects, where ***θ***_EAS_ = (0.5, 0, 0.25, 0, 0.1, 0, 0, 0, 0, 0)^*⊤*^ and ***θ***_EUR_ = (0.5, 0, 0.15, 0, 0.1, 0, 0.1, 0, 0, 0)^*⊤*^, which differed in the **3rd** and **7th** entries. That is, “fully ancestry-consistent” refers to cases where the effect sizes are identical across the two ancestries, whereas “partially ancestry-consistent” indicates that a subset of exposures have different effect sizes. We fixed the sample sizes at *n*_EUR_ = 400,000 and the number of IVs at *m*_EUR_ = 2,000 and considered *m*_EAS_ = 500 or 250 when *n*_EAS_ = 100,000 or 50,000, respectively. Likewise, we set the trait heritabilities at 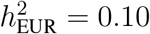 and changed 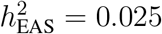 or 0.0125 when *n*_EAS_ = 100,000 or 50,000, respectively.

The overlap proportions of exposure and outcome GWAS samples are all 95%. We varied the proportion of horizontal pleiotropy to be 0%, 5%, or 10% and fixed the pleiotropy-to-exposure variance ratio at 1:2. Each scenario was repeated for 500 simulation replicates. See details in **Supplementary Materials**.

### 5.6 Stroke analysis

We applied SBayesRC (Zheng et al., 2024) to impute missing variants using reference panels containing approximately 7.3 million SNPs for EUR, 6.0 million SNPs for AFR, and 4.9 million SNPs for EAS. Next, we followed the idea of FEMA to select IVs (Hemani et al., 2025). Specifically, for each variant, we computed the *χ*^2^-statistic in combining association evidence of EUR, EAS, and AFR, and the corresponding *p*-values were calculated using a *χ*^2^-distribution with 30 degrees of freedom. In C+T, we used an alternative LD reference panel of 9,680 European individuals randomly selected from the UK Biobank. Although EAS and AFR individuals were not included, this is consistent with the principle of FEMA described by Hemani et al. (2025), which selects variants with strong effects in all ancestries. The covariance matrices of estimation errors used in the *χ*^2^ test were estimated from genome-wide SNPs with statistically insignificant associations (Zhu et al., 2015). Furthermore, IVs were clumped using PLINK (Purcell et al., 2007) with a 1 Mb window, a threshold of P*<*5E-8, and a threshold of *r*^2^*<*0.001. We further excluded the IVs that were in the loci with GEIs (Sung et al., 2018; Zhu et al., 2024). This resulted in 3,082 IVs used in multi-ancestry MVMR analysis.

In addition to the above approaches for excluding variants with potential GEIs and GGIs, we also compared two baseline strategies: (i) performing the analysis without any variant exclusion, and (ii) removing ancestry-heterogeneous variants using our proposed MR-GxE framework. Figures S2 and S3 show the results obtained without excluding variants with potential GGIs and GEIs and after removing such variants using the MR-GxE framework, respectively. For the second strategy, we performed four UVMR analyses for each exposure: using two EUR-based GWAS as exposure datasets and treating AFR and EAS GWAS as outcomes, respectively. We also conducted the inverse analyses by swapping the roles of exposure and outcome. Only genome-wide significant variants were selected as IVs, and cases with fewer than 10 IVs were excluded. All UVMR analyses were performed using MRBEE with default settings. Both sets of results are highly similar to those presented in Figure 6.

We consider TEMR as a comparator. We retained the FEMA-based IV selection strategy but applied an exposure-wise joint *χ*^2^ test with 3 degrees of freedom. For clumping and thresholding, we used `PLINK` with a 1 Mb window, a genome-wide significance threshold of P*<*5E-8, and an LD threshold of *r*^2^*<*0.001. We also excluded IVs that overlap with known GEI loci in an exposure-wise manner.

## Data and code availability

The GWAS summary data used in this study can be downloaded from the “Data available” section of the related literature. The code for the analyses presented in this paper is available in the Supplemental Materials, complete with step-by-step instructions. MRBEEX package is available at https://github.com/harryyiheyang/MRBEEX.

## Acknowledgments

This work was supported by grants HG011052 and HG011052-03S1 (to X.Z.) from the National Human Genome Research Institute (NHGRI), USA.

## Notes

https://github.com/harryyiheyang/MRBEEX

### Competing Interest Statement

The authors have declared no competing interest.

### Summary of Updates

We change the structure of the paper: Background, Results, Discussion,Methods

