## Supplementary Materials for "Improving causal effect estimation in multi-ancestry multivariable Mendelian randomization with transfer learning"

### Supplementary Materials of "Improving causal effect estimation in multi-population multivariable Mendelian randomization with transfer learning"

Yihe Yang and Xiaofeng Zhu

### 1 Supplementary Real Data Analysis

#### 1.1 Imputation using SBayesRC

Due to the scale of data, we show the quasi codes how did impute the GWAS summary data. We first merge the GWAS summary data to the reference panel provided by SBayesRC:

```
library(data.table)
library(dplyr)
library(TwoSampleMR)

variant=fread("../LDpanel/ukbEAS_Imputed/snp.info")
%>%dplyr::select(CHR=Chrom, SNP=ID, BP=PhysPos, A1, A2, Freq=A1Freq)

variant$MarkerName=paste0(variant$CHR, ":", variant$BP)
SVS=fread("../GWAS/Stroke/GCST90104548_SVS_EAS.tsv.gz")
SVS$N=effective_n(5532, 237242)
SVS=dplyr::select(SVSCHR=chromosome, BP=base_pair_location, BETA=beta,
    SE=standard_error, Freq=effect_allele_frequency,
    A1=effect_allele, A2=other_allele, P=p_value, N)

SVS$MarkerName=paste0(SVS$CHR, ":", SVS$BP)
SVS=merge(SVS, variant[, c("SNP", "MarkerName")], by="MarkerName")
```

Here, we used the effective sample size estimated by using `TwoSampleMR`, and we use `CHR:BP` as the MarkerName to merge the data. Next, we arrange the data into the COJO format and then apply SBayesRC to impute the data. Note that we did not run PRS estimation.

```
SVS=dplyr::select(SVS, SNP, A1, A2, freq=Freq, b=BETA, se=SE, p=P, N)
fwrite(SVS, "../EAS/SVS.ma", row.names=F, quote=F, sep="\t")

SBayesRC::tidy(mafile='../EAS/SVS.ma', LDdir='../LDpanel/ukbEAS_Imputed/',
 output='../EAS/SVS.ma', log2file=TRUE)

SBayesRC::impute(mafile='../EAS/SVS.ma', LDdir='../LDpanel/ukbEAS_Imputed/',
  output='../EAS/SVS.ma', log2file=TRUE)
```

For more information about SBayesRC, please visit https://github.com/zhilizheng/SBayesRC.

#### 1.2 Data Harmonization

Due to the scale of data, we show the quasi codes how did we harmonize the data. We first loaded the data:

```
library(data.table)
library(dplyr)
variant=fread("../LDpanel/ukbEAS_Imputed/snp.info")
%>%dplyr::select(CHR=Chrom, SNP=ID, BP=PhysPos, A1, A2, Freq=A1Freq)

SBP_EUR=fread("../SBP/EUR_SBP.ma")
ALB_EUR=fread("../ALB/EUR_ALB.ma")
BMI_EUR=fread("../BMI/EUR_BMI.ma")
T2D_EUR=fread("../T2D/T2D.ma")
AF_EUR=fread("../AF/EUR_AF.ma")
TG_EUR=fread("../TG/EUR_TG.ma")
SMK_EUR=fread("../SMK/EUR_SMK.ma")
LDL_EUR=fread("../LDL/EUR_LDL.ma")
```

```
HDL_EUR=fread("../HDL/EUR_HDL.ma")
AS_EUR=fread("../AS/EUR_AS.ma")
IS_EUR=fread("../IS/EUR_IS.ma")
CES_EUR=fread("../CES/EUR_CES.ma")
LAS_EUR=fread("../LAS/EUR_LAS.ma")
SVS_EUR=fread("../SVS/EUR_SVS.ma")
DRNK_EUR=fread("../DRNK/EUR_DRNK.ma")
HDL_EUR=fread("../HDL/EUR_HDL.ma")

ALB_EAS=fread("../ALB/EAS_ALB.ma")
BMI_EAS=fread("../BMI/EAS_BMI.ma")
T2D_EAS=fread("../T2D/EAS_T2D.ma")
AF_EAS=fread("../AF/EAS_AF.ma")
HDL_EAS=fread("../HDL/EAS_HDL.ma")
LDL_EAS=fread("../LDL/EAS_LDL.ma")
TG_EAS=fread("../TG/EAS_TG.ma")
SBP_EAS=fread("../SBP/EAS_SBP.ma")
SMK_EAS=fread("../SMK/EAS_SMK.ma")
AS_EAS=fread("../AS/EAS_AS.ma")
IS_EAS=fread("../IS/EAS_IS.ma")
CES_EAS=fread("../CES/EAS_CES.ma")
LAS_EAS=fread("../LAS/EAS_LAS.ma")
SVS_EAS=fread("../SVS/EAS_SVS.ma")
HDL_EAS=fread("../HDL/EAS_HDL.ma")
DRNK_EAS=fread("../DRNK/EAS_DRNK.ma")

ALB_AFR=fread("../ALB/AFR_ALB.ma")
BMI_AFR=fread("../BMI/AFR_BMI.ma")
T2D_AFR=fread("../T2D/AFR_T2D.ma")
AF_AFR=fread("../AF/AFR_AF.ma")
HDL_AFR=fread("../HDL/AFR_HDL.ma")
LDL_AFR=fread("../LDL/AFR_LDL.ma")
TG_AFR=fread("../TG/AFR_TG.ma")
SBP_AFR=fread("../SBP/AFR_SBP.ma")
SMK_AFR=fread("../SMK/AFR_SMK.ma")
AS_AFR=fread("../AS/AFR_AS.ma")
IS_AFR=fread("../IS/AFR_IS.ma")
CES_AFR=fread("../CES/AFR_CES.ma")
LAS_AFR=fread("../LAS/AFR_LAS.ma")
SVS_AFR=fread("../SVS/AFR_SVS.ma")
HDL_AFR=fread("../HDL/AFR_HDL.ma")
DRNK_AFR=fread("../DRNK/AFR_DRNK.ma")
```

Note that after SBayesRC's imputation, the data were automatically allele harmonized. Therefore, we could directly combine them:

```
GWAS_EUR=list(AF=AF_EUR, ALB=ALB_EUR, BMI=BMI_EUR, DRNK=DRNK_EUR, `HDL-C`=HDL_EUR,
    `LDL-C`=LDL_EUR, SBP=SBP_EUR, SMK=SMK_EUR, T2D=T2D_EUR, TG=TG_EUR,
    AS=AS_EUR, IS=IS_EUR, LAS=LAS_EUR, CES=CES_EUR, SVS=SVS_EUR)

GWAS_EAS=list(AF=AF_EAS, ALB=ALB_EAS, BMI=BMI_EAS, DRNK=DRNK_EAS, `HDL-C`=HDL_EAS,
    `LDL-C`=LDL_EAS, SBP=SBP_EAS, SMK=SMK_EAS, T2D=T2D_EAS, TG=TG_EAS,
    AS=AS_EAS, IS=IS_EAS, LAS=LAS_EAS, CES=CES_EAS, SVS=SVS_EAS)
```

```
GWAS_AFR=list(AF=AF_AFR, ALB=ALB_AFR, BMI=BMI_AFR, DRNK=DRNK_AFR, `HDL-C`=HDL_AFR,
    `LDL-C`=LDL_AFR, SBP=SBP_AFR, SMK=SMK_AFR, T2D=T2D_AFR, TG=TG_AFR,
    AS=AS_AFR, IS=IS_AFR, LAS=LAS_AFR, CES=CES_AFR, SVS=SVS_AFR)

Z_EUR=N_EUR=matrix(0, 7356518, 15)
Z_EAS=N_EAS=matrix(0, 4906538, 15)
Z_AFR=N_AFR=matrix(0, 6064174, 15)
for(i in 1:15){
Z_EUR[, i]=GWAS_EUR[[i]]$b/GWAS_EUR[[i]]$se
N_EUR[, i]=GWAS_EUR[[i]]$N
Z_EAS[, i]=GWAS_EAS[[i]]$b/GWAS_EAS[[i]]$se
N_EAS[, i]=GWAS_EAS[[i]]$N
Z_AFR[, i]=GWAS_AFR[[i]]$b/GWAS_AFR[[i]]$se
N_AFR[, i]=GWAS_AFR[[i]]$N
}
rownames(Z_EUR)=rownames(N_EUR)=GWAS_EUR$ALB$SNP
rownames(Z_EAS)=rownames(N_EAS)=GWAS_EAS$ALB$SNP
rownames(Z_AFR)=rownames(N_AFR)=GWAS_AFR$ALB$SNP
colnames(Z_EUR)=colnames(Z_EAS)=colnames(N_EUR)=colnames(N_EAS)=names(GWAS_EAS)
colnames(Z_AFR)=colnames(Z_AFR)=names(GWAS_AFR)
SNPOverlap=intersect(GWAS_AFR$AF$SNP, GWAS_EAS$AF$SNP)
Z_EUR=Z_EUR[SNPOverlap, ];N_EUR=N_EUR[SNPOverlap, ];
Z_AFR=Z_AFR[SNPOverlap, ];N_AFR=N_AFR[SNPOverlap, ];
Z_EAS=Z_EAS[SNPOverlap, ];N_EAS=N_EAS[SNPOverlap, ];
```

According to the principle of FEMA, we would use the set of IVs that were strongly associated in both EUR and EAS, and hence here we would exclude the variants that were not in the LD reference panel of EAS (number of variants is 4, 906, 538).

Next, we estimate the correlation matrix of estimation errors for bias-correction:

```
Rxy_EUR=MRBEE::errorCov(ZMatrix=Z_EUR, subsampling.ratio=0.05, subsampling.time=300, Zscore.cutoff=2.2)
Rxy_EAS=MRBEE::errorCov(ZMatrix=Z_EAS, subsampling.ratio=0.05, subsampling.time=300, Zscore.cutoff=2.2)
Rxy_AFR=MRBEE::errorCov(ZMatrix=Z_AFR, subsampling.ratio=0.05, subsampling.time=300, Zscore.cutoff=2.2)
colnames(Rxy_EUR)=rownames(Rxy_EUR)=colnames(Z_EUR)
colnames(Rxy_EAS)=rownames(Rxy_EAS)=colnames(Z_EUR)
colnames(Rxy_AFR)=rownames(Rxy_AFR)=colnames(Z_EUR)
Theta_EUR=solve(Rxy_EUR[1:10, 1:10])
Theta_EAS=solve(Rxy_EAS[1:10, 1:10])
Theta_AFR=solve(Rxy_AFR[1:10, 1:10])
```

We found to set `Zscore.cutoff` slightly larger than 2 could be better.

#### 1.3 Fixed-Effect Meta-Analysis

Next, we selected the IVs according to fixed-effect meta-analysis (FEMA). We first calculated the $\chi^2$-statistics and yielded the $p$-values:

```
Theta_EUR=solve(Rxy_EUR[1:10, 1:10])
Theta_EAS=solve(Rxy_EAS[1:10, 1:10])
PV=c(1:nrow(Z_EUR))
for(i in 1:length(PV1)){
z1=Z_EUR[i, 1:10]
z2=Z_EAS[i, 1:00]
z3=Z_AFR[i, 1:00]
```

```
chi1=sum(z1*(Theta_EUR%*%z1))
chi2=sum(z2*(Theta_EAS%*%z2))
chi3=sum(z3*(Theta_AFR%*%z3))
pv=pchisq(chi1+chi2, 30, lower.tail=F)
PV[i]=pv
}
```

We then extracted the IVs using PLINK:

```
library(data.table)
library(dplyr)
library(arrow)
A=data.frame(SNP=rownames(Z_EUR), P=PV)%>%filter(P<5e-7)
fwrite(A, "../plinkfile/A.txt", row.names=F, quote=F, sep="\t")

system("./plink --bfile ../NealeReference/all_chrs --clump ../plinkfile/A.txt
--clump-field P  --clump-kb 1000 --clump-p1 2.5e-09 --clump-p2 2.5e-09
--clump-r2 0.01  --out  ../plinkfile/A")

plinkfile=fread("../plinkfile/A.clumped")
  %>%dplyr::select(SNP, CHR, BP, P)

Z_EUR=Z_EUR[plinkfile$SNP, ]
N_EUR=N_EUR[plinkfile$SNP, ]
Z_EAS=Z_EAS[plinkfile$SNP, ]
N_EAS=N_EAS[plinkfile$SNP, ]
Z_AFR=Z_AFR[plinkfile$SNP, ]
N_AFR=N_AFR[plinkfile$SNP, ]
saveRDS(list(Z_EUR=Z_EUR, Z_EAS=Z_EAS, Z_AFR=Z_AFR, N_EUR=N_EUR, N_AFR=N_AFR,
   N_EAS=N_EAS, Rxy_EUR=Rxy_EUR, Rxy_AFR=Rxy_AFR, Rxy_EAS=Rxy_EAS),
   "../Clumping.rds")
```

#### 1.4 Removing IVs with GEIs

We found a vector of variants that had genome-wide significant GEI effects, which were extracted from Zhu et al. [10] (Table S3) and Sung et al. [7] (Table S10). We first load the data:

```
Clumping=readRDS("RDS/Clumping.rds")
SNPList=rownames(Clumping$Z_EUR)
V=read_parquet("RDS/7M_with_Freq.parquet")
V=V[which(V$SNP%in%SNPList), ]
GxESNP=readRDS("RDS/GxESNP.rds")
```

We then excluded the variants that were in the +/- 1MB window of the a GEI signals.

```
V=as.data.table(V)
GxESNP=as.data.table(GxESNP)
to_remove_idx=logical(nrow(V))

for (i in 1:nrow(GxESNP)) {
chr_i=GxESNP$CHR[i]
bp_i=GxESNP$BP[i]
bp_min=bp_i-1e6
bp_max=bp_i+1e6
to_remove_idx=to_remove_idx|(V$CHR==chr_i&V$BP>= bp_min&V$BP<=bp_max)
```

```
}

V_filtered=V[!to_remove_idx, ]
SNP_to_keep=V_filtered$SNP
ind=which(V$SNP%in%SNP_to_keep)

Clumping$Z_EUR=Clumping$Z_EUR[ind, ]
Clumping$Z_EAS=Clumping$Z_EAS[ind, ]
Clumping$Z_AFR=Clumping$Z_AFR[ind, ]
Clumping$N_EUR=Clumping$N_EUR[ind, ]
Clumping$N_EAS=Clumping$N_EAS[ind, ]
Clumping$N_AFR=Clumping$N_AFR[ind, ]
saveRDS(Clumping, "RDS/Clumping_Excluded.rds")
```

##### 1.5 Analysis for all strokes

We present the analysis for all strokes as an example, noting that analyses for other stroke subtypes require only minor modifications. The provided dataset includes GWAS summary statistics for all five stroke subtypes. We first performed MRBEE on the EUR data to obtain source causal effect estimates:

```
library(data.table)
library(dplyr)
library(arrow)
library(MRBEE)
library(MRBEEX)
readRDS("RDS/Clumping_Excluded.rds") %>% list2env(., envir=.GlobalEnv)
```

```
## <environment: R_GlobalEnv>
```

```
BETA=Z_EUR/sqrt(N_EUR)
SE=1/sqrt(N_EUR)
bX=BETA[, 1:10]
bXse=SE[, 1:10]
by=BETA[, 11]
byse=SE[, 11]
NAM=c(1:10, 11)
fit_EUR=MRBEE_TL(by, bX, byse, bXse, Rxy=Rxy_EAS[NAM, NAM], theta.source=bX[1, ]*0,
       theta.source.cov=diag(bX[1, ])*0, pip.thres=0.5, ebic.delta=0,
       ebic.gamma=2, admm.rho=3, Lvec=c(1:10), sampling.time=1000,
       tauvec=seq(5, 20, 1), reliability.thres=0.8, pip.min=0.2)
```

```
## Bootstrapping process:
##   |                                                                   |
```

```
theta.source=fit_EUR$theta;
theta.source.cov=fit_EUR$theta.cov;
```

By letting `theta.source=bX[1, ]*0` and `theta.source.cov=diag(bX[1, ])*0`, we can impose sparsity on `fit_EUR$theta`.

We also applied MRBEE on the EAS data specifically:

```
BETA=Z_EAS/sqrt(N_EAS)
SE=1/sqrt(N_EAS)
bX=BETA[, 1:10]
bXse=SE[, 1:10]
```

```r
by=BETA[, 11]
byse=SE[, 11]
NAM=c(1:10, 11)
fit_EAS=MRBEE_IMRP(by=by, bX=bX, byse=byse, bXse=bXse, Rxy=Rxy_EAS[NAM, NAM], var.est="variance")
```

Next, we implemented MRBEE-TL with $\hat{\boldsymbol{\theta}}_{\text{source}}$ and $\text{cov}(\hat{\boldsymbol{\theta}}_{\text{source}})$ (this will be used in SE estimation):

```r
fit_EAS_Tran=MRBEE_TL(by, bX, byse, bXse, Rxy=Rxy_EAS[NAM, NAM],
          theta.source=theta.source, theta.source.cov=theta.source.cov,
          transfer.coef=1, pip.thres=0.5, LD="identity",
          ebic.delta=0, ebic.gamma=2, admm.rho=3, Lvec=c(1:10),
          sampling.time=1000, tauvec=seq(5, 20, 1))
```

```
## Bootstrapping process:
##   |                                                                      |
```

```r
BETA=Z_AFR/sqrt(N_AFR)
SE=1/sqrt(N_AFR)
bX=BETA[, 1:10]
bXse=SE[, 1:10]
by=BETA[, 11]
byse=SE[, 11]
NAM=c(1:10, 11)
fit_AFR=MRBEE_IMRP(by=by, bX=bX, byse=byse, bXse=bXse, Rxy=Rxy_AFR[NAM, NAM], var.est="variance")
fit_AFR$theta/fit_AFR$theta.se
```

```
##         AF        ALB        BMI       DRNK      HDL-C      LDL-C        SBP
##  2.3309864 -1.7997874 -1.6738018 -0.1093087 -1.6478910  2.2448588  1.6873797
##        SMK        T2D         TG
##  1.1093570  2.6192736 -1.2268205
```

```r
fit_AFR_Tran=MRBEE_TL(by, bX, byse, bXse, Rxy=Rxy_AFR[NAM, NAM], theta.source=theta.source,
           theta.source.cov=theta.source.cov, pip.thres=0.5,
           ebic.delta=0, ebic.gamma=2, admm.rho=3, Lvec=c(1:5),
           sampling.time=1000, tauvec=seq(5, 20, 1))
```

```
## Bootstrapping process:
##   |                                                                      |
```

```r
library(ggplot2)
library(data.table)
Estimate=cbind(fit_EUR$theta, fit_EAS_Tran$theta, fit_EAS$theta, fit_AFR_Tran$theta, fit_AFR$theta)
Estimate
```

```
##              [,1]        [,2]        [,3]        [,4]         [,5]
## AF     0.17064367  0.04735572  0.06519989  0.07974070  0.166525479
## ALB   -0.07405796 -0.07405796 -0.03057905 -0.07405796 -0.166752594
## BMI    0.00000000  0.00000000 -0.03755217  0.00000000 -0.097882714
## DRNK   0.00000000  0.07918363  0.10096425  0.00000000 -0.009441188
## HDL-C  0.00000000  0.00000000  0.01133246  0.00000000 -0.101664674
## LDL-C  0.05363107  0.05363107  0.04965362  0.05363107  0.207563989
## SBP    0.24608986  0.50606782  0.52293391  0.12905798  0.123636747
## SMK    0.09895466  0.09895466  0.04447729  0.09895466  0.141864465
## T2D    0.09242643  0.00000000  0.02280425  0.09242643  0.144746721
```

```
## TG     0.00000000  0.00000000  0.01381294  0.00000000 -0.109003418
```

```
SE=cbind(fit_EUR$theta.se, fit_EAS_Tran$theta.se, fit_EAS$theta.se, fit_AFR_Tran$theta.se, fit_AFR$theta
Z=Estimate/SE
colnames(Z)=colnames(Estimate)=c("European (specific)", "East Asian (transfer learning)",
                                 "East Asian (specific)", "African American (transfer learning)",
                                 "African American (specific)")
Estimate=as.data.frame(as.table(Estimate))
colnames(Estimate)=c("Exposure", "Population", "Estimate")
SE=as.data.frame(as.table(SE))
colnames(SE)=c("Exposure", "Population", "SE")
G=Estimate
G$SE=SE$SE
G$P=pchisq((G$Estimate/G$SE)^2, 1, lower.tail=F)
G$Outcome="All Strokes (AS)"
G$Annotation1=0
G$Annotation1[c(2, 12, 6, 16, 8, 18)]=1
G$Annotation2=0
G$Annotation2[c(2, 32, 6, 36, 8, 38, 9, 39)]=1

trait_table=data.table(
Abbreviation=c("AF", "ALB", "BMI", "DRNK", "HDL-C", "LDL-C", "SBP", "SMK", "T2D", "TG"),
FullName=c(
"Atrial Fibrillation",
"Serum Albumin",
"Body Mass Index",
"Drinks Per Week",
"HDL Cholesterol",
"LDL Cholesterol",
"Systolic Blood Pressure",
"Smoking Initiation",
"Type 2 Diabetes",
"Triglycerides"
)
)
G$lower=G$Estimate - G$SE * 2
G$upper=G$Estimate + G$SE * 2
G$Outcome=ordered(G$Outcome, levels=c("All Strokes (AS)",
                              "All Ischemic Strokes (IS)",
                              "Large-Artery Atherosclerotic Stroke (LAS)",
                              "Cardioembolic Stroke (CES)",
                              "Small Vessel Stroke (SVS)"))
G$Significant=ifelse(G$lower * G$upper <= 0, "No", "Yes")
G$EAS_Transferred=ifelse(G$Annotation1==1, "Yes", "No")
G$AFR_Transferred=ifelse(G$Annotation2==1, "Yes", "No")
G$Transfer_Label=NA_character_
G$Transfer_Label[G$Annotation1==1&G$Annotation2==1]="EEA"
G$Transfer_Label[G$Annotation1==1&G$Annotation2 != 1]="EE"
G$Transfer_Label[G$Annotation1 != 1&G$Annotation2==1]="EA"
target_populations=c(
"European (specific)",
"East Asian (transfer learning)",
"African American (transfer learning)"
)
```

```
dt_eur_eea=unique(G[G$Population=="European (specific)"&
                      G$Transfer_Label=="EEA", c("Exposure", "Outcome")])
for (i in seq_len(nrow(dt_eur_eea))) {
idx <- G$Exposure == dt_eur_eea$Exposure[i] &
G$Outcome == dt_eur_eea$Outcome[i] &
G$Population %in% target_populations
G$Transfer_Label[idx] <- "EEA"
}
G$star_y <- ifelse(G$Estimate >= 0, G$upper + 0.1, G$lower - 0.1)
G$Population <- ordered(
G$Population,
levels=rev(c(
"African American (specific)",
"East Asian (specific)",
"African American (transfer learning)",
"East Asian (transfer learning)",
"European (specific)"
))
)
G$Exposure <- factor(G$Exposure, levels=unique(G$Exposure))

ggplot(G, aes(x=Exposure, y=Estimate, fill=Population)) +
geom_bar(stat="identity", position=position_dodge(width=0.8), width=0.6, color="black") +
geom_errorbar(
aes(ymin=lower, ymax=upper),
position=position_dodge(width=0.8), width=0.2) +
geom_text(
aes(y=star_y, label=Transfer_Label),
position=position_dodge(width=0.8),
vjust=0.75, size=2.5, hjust=0.5,
na.rm=TRUE) +
facet_wrap(Outcome ~ ., nrow=5, scales="free_y") +
scale_fill_manual(
values=c(
"European (specific)"="#ee2560",
"East Asian (transfer learning)"="#feee7d",
"African American (transfer learning)"="#45d9fd",
"East Asian (specific)"="#f9c00c",
"African American (specific)"="#2b90d9"
),
name="Population") +
geom_vline(xintercept=0, linetype="dashed") +
theme_bw() +
theme(
strip.text=element_text(size=16),
axis.text.x=element_text(hjust=0.5, size=10, angle=0),
axis.text.y=element_text(size=12),
axis.title=element_text(size=12),
legend.position="bottom",
legend.text=element_text(size=12),
legend.title=element_blank(),
panel.spacing=unit(1.2, "lines"),
panel.grid.major.y=element_blank(),
```

```
panel.grid.major.x=element_line(color="grey80", linewidth=0.5),
plot.caption=element_text(hjust=0.5, size=11, margin=margin(t=12))
) +
labs(
x=NULL,
y="causal effect estimate",
caption="EEA: consistent in EUR, EAS, and AFA        EE: consistent in EUR and EAS        EA: consistent i
)
```

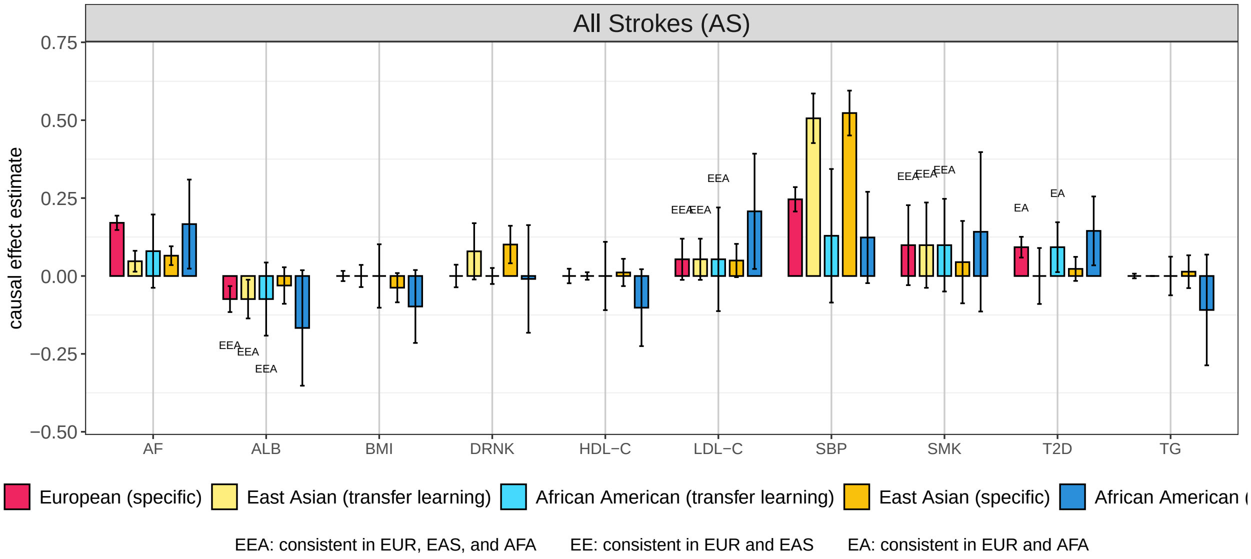

### 2 Supplemetary Simulations

#### 2.1 Simulation Settings

We first illustrate the settings of our simulation. The following part of codes demonstrate the basic settings including the sample sizes, the number of IVs, the genetic correlation matrix of exposures, etc.

```
library(MASS)
library(MendelianRandomization)
library(MRBEEX)
ARcov=function(p, rho){
s=c(1:p)
for(i in 1:p){
s[i]=rho^(i-1)
}
return(toeplitz(s))
}
CScov=function(p, rho){
A=matrix(rho, p, p)+(1-rho)*diag(p)
return(A)
}
# two structured covariance matrix function
theta_eur=c(1, 0, 0.3, 0, 0.2, 0, 0.2, rep(0, 3))/2
theta_eas=c(1, 0, 0.5, 0, 0.2, 0, 0, rep(0, 3))/2
# theta_eas=theta_eur
# Implement this for all ancestry-consistent case
n_eas=1e5/2
n_eur=4e5
m_eas=500/2
m_eur=2000
h_eur=0.1
h_eas=0.025/2
SB_eur=kronecker(CScov(2, 0.5), ARcov(5, 0.5))
SB_eas=kronecker(CScov(2, 0.6), ARcov(5, 0.4))
# We consider slightly different genetic correlation of exposures
SE_eur=SE_eas=CScov(11, 0.3)
# The correlation matrix of estimation errors are fixed for both population
Rnn=CScov(11, 0.95)
# Sample-overlap proportion matrix
UHP.frac=0.05*2
UHP.var=0.5
```

#### 2.2 Simulation Implementation

The next part of codes generate the GWAS summary data, by using `summary_generation` provided by `MRBEEX`.

```
A_eur=summary_generation(theta=theta_eur, m=m_eur, Rbb=SB_eur, Ruv=SE_eur, Rnn=Rnn,
         LD="identity", Nxy=rep(n_eur, 11), non.zero.frac=rep(0.8, 10),
         UHP.frac=UHP.frac, CHP.frac=0, UHP.var=UHP.var,
         Hxy=rep(h_eur, 11), UHP.dis="uniform")
A_eas=summary_generation(theta=theta_eas, m=m_eas, Rbb=SB_eas, Ruv=SE_eas, Rnn=Rnn,
         LD="identity", Nxy=rep(n_eas, 11), non.zero.frac=rep(0.8, 10),
         UHP.frac=UHP.frac, CHP.frac=0, UHP.var=UHP.var,
         Hxy=rep(h_eas, 11), UHP.dis="uniform")
```

```
bX_eur=A_eur$bX
by_eur=A_eur$by
bXse_eur=A_eur$bXse
byse_eur=A_eur$byse
Rxy_eur=A_eur$Rxy
bX_eas=A_eas$bX
by_eas=A_eas$by
bXse_eas=A_eas$bXse
byse_eas=A_eas$byse
Rxy_eas=A_eas$Rxy
colSums(A_eur$bX0^2)
```

```
##  [1] 0.1 0.1 0.1 0.1 0.1 0.1 0.1 0.1 0.1 0.1
```

```
# Our function fixes the heritability of exposures to the given values
sum(A_eur$by0^2/A_eur$Syy)
```

```
## [1] 0.09747686
```

```
# For outcome, it also fixes the heritability
# This function only fix var(X)=1 and allow var(y)>1.
# Rxy is also not a correlation matrix
# This may be okay in simulation
```

Finally, we reach the part of codes implementing each method:

```
fit_eur_mrbee=MRBEE_IMRP(by=by_eur, bX=bX_eur, byse=byse_eur, bXse=bXse_eur, Rxy=Rxy_eur)
theta.source=fit_eur_mrbee$theta
ind=which(abs(theta.source/fit_eur_mrbee$theta.se)<1.96)
theta.source[ind]=0
theta.source.cov=fit_eur_mrbee$covtheta
theta.source.cov[ind, ind]=0
# get the source data
t1=Sys.time()
fit_mrbee=MRBEE_IMRP(by=by_eas, bX=bX_eas, byse=byse_eas, bXse=bXse_eas, Rxy=Rxy_eas)
t2=Sys.time()
imrp.time=difftime(t2, t1, units="secs")

MVINPUT=mr_mvinput(by=by_eas, bx=bX_eas, byse=byse_eas, bxse=bXse_eas)
t1=Sys.time()
fit_median=mr_mvmedian(MVINPUT, iterations=1000)
t2=Sys.time()
median.time=difftime(t2, t1, units="secs")

t1=Sys.time()
fit_lasso=mr_mvlasso(MVINPUT)
t2=Sys.time()
lasso.time=difftime(t2, t1, units="secs")

t1=Sys.time()
fit_mvcML=mr_mvcML(MVINPUT, n=n_eas, DP=F, rho_mat=Rxy_eas)
t2=Sys.time()
mvcML.time=difftime(t2, t1, units="secs")

t1=Sys.time()
fit_tran_mrbee=MRBEE_TL(by=by_eas, bX=bX_eas, byse=byse_eas, bXse=bXse_eas,
```

```
        Rxy=Rxy_eas, theta.source=theta.source,
        theta.source.cov=theta.source.cov)
```

```
## Bootstrapping process:
##   |                                                                              |
```

```
t2=Sys.time()
mrbee.tran.time=difftime(t2, t1, units="secs")

Estimate=cbind(fit_tran_mrbee$theta, fit_mrbee$theta,
fit_mvcML@Estimate, fit_median@Estimate, fit_lasso@Estimate)
colnames(Estimate)=c("MRBEE-TL", "MRBEE", "MRcML", "MRMedian", "MRLasso")
time=c(mrbee.tran.time, imrp.time, mvcML.time, median.time, lasso.time)
names(time)=c("MRBEE-TL", "MRBEE", "MRcML", "MRMedian", "MRLasso")
print(apply(Estimate-theta_eas, 2, norm, "2"))
```

```
##  MRBEE-TL     MRBEE     MRcML  MRMedian   MRLasso
## 0.1412880 0.2164963 0.2577930 0.1906517 0.2522305
```

```
print(time)
```

```
## Time differences in secs
##   MRBEE-TL      MRBEE      MRcML   MRMedian    MRLasso
## 1.48344707 0.00732708 7.96975708 0.68373609 0.06393194
```

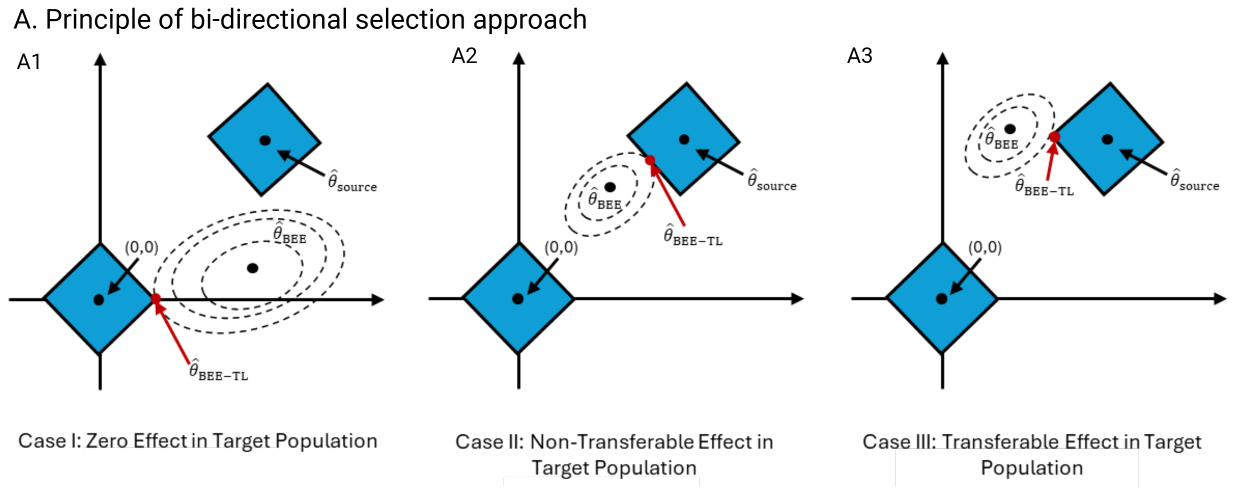

Figure S1: Illustration of principle of BDSP.

### 3 Supplementary Method

In this section, we give the detailed algorithm and related implementation issues of MRBEE-TL.

#### 3.1 Illustration of bidirectional separation penalty

We first illustrate the principle of bidirectional separation penalty (BDSP) by using the panels A1 , A2, and A3 in Figure S1, which is Figure 1A in the main text.

To visualize how MRBEE-TL performs estimation under different causal scenarios, Figure S1 illustrates three representative cases in a simplified setting with $p = 2$ exposures. The loss function in MRBEE-TL combines a bias-corrected estimating equation loss, which yields elliptical contours centered at $\hat{\boldsymbol{\theta}}_{\text{BEE}}$, and a BDSP, which introduces two competing $\ell_1$-type constraints centered at $(0, 0)$ and $\hat{\boldsymbol{\theta}}_{\text{source}}$, respectively.

In this illustration, we consider introduce BDSP by using lasso, but it can be extended to other penalty such as MCP and SuSiE. Specifically, it is formally defined as:

$$\rho_\lambda(\boldsymbol{\theta}; \mathbf{0}, \hat{\boldsymbol{\theta}}_{\text{source}}) = \lambda \sum_{j=1}^{p} \min\left\{ d(\theta_j, 0), d(\theta_j, \hat{\theta}_{\text{source},j}) \right\}, \tag{1}$$

where $d(x_1, x_2)$ is a distance, which can be the $\ell_1$-norm

$$d(x_1, x_2) = |x_1 - x_2|,$$

and can also be the $\ell_2$-norm

$$d(x_1, x_2) = d(x_1 - x_2) = \frac{1}{2}(x_1 - x_2)^2.$$

This penalty encourages each entry $\theta_j$ to be either zero (sparsity) or similar to the corresponding source estimate $\hat{\theta}_{\text{source},j}$ (consistency).

Figure S1 overlays the elliptical contours of the MRBEE loss with two diamond-shaped constraint regions corresponding to the $\ell_1$ penalties in BDSP. The MRBEE-TL estimate $\hat{\boldsymbol{\theta}}_{\text{BEE-TL}}$ is obtained as the first point of intersection between the elliptical contours and the union of the two diamonds. This is analogous to the classical Lasso interpretation, where the solution lies on the boundary of an $\ell_1$ constraint.

- **Case I (Panel A1): Zero Effect in Target Population.** In this setting, the true target effect is zero. The elliptical contours touch the diamond centered at $(0, 0)$ first, yielding:

$$\hat{\boldsymbol{\theta}}_{\text{BEE-TL}} = (r, 0), \tag{2}$$

where $r$ is the radius of the diamond, which means correctly identifying the absence of causal effects in the target population.

- **Case II (Panel A2): Ancestry-inconsistent Effect.** Here, the target effect differs from both zero and the source effect. The loss contour touches the boundary of the source estimate constraint (but not at a corner), and the solution remains close to the unconstrained MRBEE estimate:

$$\hat{\boldsymbol{\theta}}_{\text{BEE-TL}} \approx \hat{\boldsymbol{\theta}}_{\text{BEE}}, \tag{3}$$

but may suffer from small bias when lasso is used. We refer to this as an ancestry-specific effect.

- **Case III (Panel A3): Ancestry-consistent Effect.** The true effect in the target population is similar to the source. The contour first intersects the diamond centered at $\hat{\boldsymbol{\theta}}_{\text{source}}$ at a corner, resulting in:

$$\hat{\boldsymbol{\theta}}_{\text{BEE-TL}} = (\hat{\boldsymbol{\theta}}_{\text{source, 1}} - r, \hat{\boldsymbol{\theta}}_{\text{source, 2}}), \tag{4}$$

which is a sign of successfully transferring the source effect to the target population for the second coordinate.

This geometric interpretation demonstrates how MRBEE-TL balances fidelity to the data (via elliptical contours) and prior structural assumptions (via BDSP), enabling robust inference even under heterogeneity. For a detailed explanation of this principle, see **hastie2015statistical** (Figure 2.2) and Tang, Xue, and Qu [8] (Figure 1 and Figure 2).

#### 3.2 Equivalence between bidirectional separation penalty and clustering

We demonstrate that the bidirectional separation penalty, as implemented in center-based regularization frameworks [4], is equivalent to a clustering procedure. Consider $n$ individuals with regression coefficients $\boldsymbol{\beta}_1, \ldots, \boldsymbol{\beta}_n$, and suppose there exist $K$ latent clusters characterized by unknown group centers $\boldsymbol{\gamma}_1, \ldots, \boldsymbol{\gamma}_K$. The center-augmented objective function is defined as

$$S(\boldsymbol{\beta}) = \frac{1}{2n}\sum_{i=1}^{n}(y_i - \boldsymbol{x}_i^\top \boldsymbol{\beta})^2 + \lambda \sum_{j=1}^{p} \min_{k=1,\ldots,K} d(\beta_j - \gamma_{kj}), \tag{5}$$

where $\lambda > 0$ is a tuning parameter, and the penalty term enforces proximity of each $\boldsymbol{\beta}_i$ to its nearest center. The penalty

$$\rho_\lambda(\beta, \gamma_1, \ldots, \gamma_k) = \lambda \sum_{j=1}^{p} \min_{k=1,\ldots,K} d(\beta - \gamma_k) \tag{6}$$

is called the multidirectional separation penalty (MDSP) where BDSP is a specific case of it (dimension = 2).

We define a latent membership indicator $\delta_j \in \{1, \ldots, K\}$ such that

$$\delta_j = \arg\min_k |\beta_j - \gamma_{kj}|. \tag{7}$$

Then the penalty term can be equivalently written as

$$\lambda \sum_{i=1}^{n} \tilde{d}(\delta_j), \tag{8}$$

where

$$\tilde{d}(x_1 - x_2) = d(x_1, x_2)$$

is a univariable function of the distance.

This establishes that the bidirectional center-based penalty is not merely a regularization device, but a computationally efficient surrogate for subgroup identification via clustering:

$$\text{penalty: } \sum_{j=1}^{p} \min_k d(\boldsymbol{\beta}_j, \boldsymbol{\gamma}_{jk}) \quad \Longleftrightarrow \quad \text{cluster assignment of } \boldsymbol{\beta}_j. \tag{9}$$

In the MRBEE-TL framework, a special case of $K = 2$ arises when we consider two competing centers: $\gamma_1 = 0$ for the zero (non-causal) group, and $\gamma_2 = \hat{\theta}_{\text{source}}$ representing the ancestry-consistent causal effect from the source population. The bidirectional separation penalty

$$\rho_\lambda(\theta_j; 0, \hat{\theta}_{\text{source},j}) = \min\left(\tilde{\rho}_\lambda(\theta_j), \tilde{\rho}_\lambda(\theta_j - \hat{\theta}_{\text{source},j})\right) \tag{10}$$

then corresponds to clustering each $\theta_j$ toward either 0 or $\hat{\theta}_{\text{source},j}$, depending on which penalty yields better model fit, mirroring a decision between "non-causal" and "ancestry-consistent causal" states.

##### 3.3 Update of causal effect estimate

We first work with BDSP (equation 11 in the main body):

$$\begin{aligned}\boldsymbol{\delta}^{(t+1)} = \arg\min_{\boldsymbol{\theta}} \Bigg\{ & \frac{1}{2m} \sum_{j=1}^{m} (\hat{\alpha}_j - \hat{\boldsymbol{\beta}}_j^\top \boldsymbol{\vartheta}^{(t+1)} - \gamma_j^{(t)} - \hat{\boldsymbol{\beta}}_j^\top \boldsymbol{\delta})^2 - \frac{1}{2} \boldsymbol{\delta}^\top \boldsymbol{\Sigma}_{W_\beta W_\beta} \boldsymbol{\delta} \\ & + (\boldsymbol{\sigma}_{W_\beta w_\alpha} - \boldsymbol{\Sigma}_{W_\beta W_\beta} \boldsymbol{\vartheta}^{(t+1)})^\top \boldsymbol{\delta} + \sum_{k=1}^{p} \tilde{\rho}_\lambda(\delta_k) \Bigg\}. \end{aligned} \tag{11}$$

Here, $\tilde{\rho}_\lambda(\cdot)$ is a penalty function with a vector of involved parameter $\lambda$. Traditional variable selection penalties include lasso [**tibshirani1996regression**] and nonconvex penalties, such as those in the class encompassing smoothly clipped absolute deviation (SCAD) [**fan2001variable**] and MCP [**zhang2010nearly**]. On the other hand, from the Bayesian point of view, $\tilde{\rho}_\lambda(x) \propto -\log f_\lambda(x)$ where $f_\lambda(x)$ is the density function of the prior distribution of a random variable $x$. **bondell2012consistent**, **moreno2015posterior**, and **tang2018bayesian** has shown that many Bayesian priors, including the global-local shrinkage priors and spiked-and-slab priors, can be viewed as special cases of nonconvex penalties.

In MRBEE-TL, SuSiE is used as $\tilde{\rho}_\lambda(\cdot)$ which is the most popular fine-mapping method for the summarized-statistic-based model. Specifically, as described in Wang et al. [9], SuSiE considers the following hierarchical Bayesian model:

$$\begin{aligned} & y = \boldsymbol{X}\boldsymbol{\beta} + \boldsymbol{e}, \quad \boldsymbol{e} \sim \mathcal{N}(0, \sigma^2 I_n), \\ & \boldsymbol{\beta} = \sum_{l=1}^{L} \mathbf{b}_l, \quad \mathbf{b}_l \sim \gamma_l b_l, \\ & \gamma_l \sim \text{Mult}(1, \pi), \quad b_l \sim \mathcal{N}_p(0, \sigma^2 \sigma_{0l}^2), \end{aligned} \tag{12}$$

It is challenging to explicitly specify the prior for the parameter $\boldsymbol{\beta}$. However, Wang et al. [9] demonstrate in their supplementary materials that, for a fixed $L$, as the number of variables $p \to \infty$, SuSiE is equivalent to Bayesian variable selection regression (BVSR) based on a spiked-and-slab prior.

We use the function `susie_suff_stat()` to update (11). Specifically, this function requires to input `XtX`,

Xty, and yty, which are respectively:

$$\text{XtX} = \sum_{j=1}^{m} \hat{\boldsymbol{\beta}}_j \hat{\boldsymbol{\beta}}_j^\top - m^* \boldsymbol{\Sigma}_{W_\beta W_\beta}, \tag{13}$$

$$\text{Xty} = \sum_{j=1}^{m} (\hat{\alpha}_j - \hat{\boldsymbol{\beta}}_j^\top \boldsymbol{\vartheta}^{(t+1)} - \gamma_j^{(t)}) \hat{\boldsymbol{\beta}}_j - m^* (\boldsymbol{\sigma}_{W_\beta w_\alpha} - \boldsymbol{\Sigma}_{W_\beta W_\beta} \boldsymbol{\vartheta}^{(t+1)}), \tag{14}$$

$$\text{yty} = \sum_{j=1}^{m} (\hat{\alpha}_j - \hat{\boldsymbol{\beta}}_j^\top \boldsymbol{\vartheta}^{(t+1)} - \gamma_j^{(t)})^2, \tag{15}$$

where $m^*$ is the number of valid IVs, i.e., the number of zero entries in $\boldsymbol{\gamma}^{(t)}$. Sample size is $m$ in `susie_suff_stat()` and $L$ will be selected by EBIC [**chen2008extended**] (described latter).

We notice that the direct output of SuSiE software is a little bias. This is due to that it also incorporates a prior distribution $b_l \sim \mathcal{N}_p(0, \sigma^2 \sigma_{0l}^2)$ which is equivalent to the ridge penalty and hence there is a shrinkage bias. To solve this problem, we set a augment `pip.thres`, where the posterior inclusion probabilities (PIPs) of entries in $\boldsymbol{\delta}^{(t+1)}$ larger than this threshold will be considered to be selected by SuSiE. Let $\mathcal{M}_\theta^{(t)}$ be such a index set of these non-zero entries. We then apply the original MRBEE to reestimate these entries by:

$$\boldsymbol{\delta}_{\mathcal{M}_\theta^{(t)}}^{(t+1)} = \text{XtX}_{\mathcal{M}_\theta^{(t)} \mathcal{M}_\theta^{(t)}}^{-1} \text{Xty}_{\mathcal{M}_\theta^{(t)}}, \tag{16}$$

where $\text{XtX}_{\mathcal{M}_\theta^{(t)} \mathcal{M}_\theta^{(t)}}$ is a submatrix of XtX with columns and rows in $\mathcal{M}_\theta^{(t)}$, and $\boldsymbol{\delta}_{\mathcal{M}_\theta^{(t)}}^{(t+1)}$ and $\text{Xty}_{\mathcal{M}_\theta^{(t)}}$ are two subvectors of $\boldsymbol{\delta}^{(t+1)}$ and Xty with entries in $\mathcal{M}_\theta^{(t)}$.

#### 3.4 Update of horizontal pleiotropy

The minimization for updating $\boldsymbol{\gamma}$ is

$$\boldsymbol{\gamma}^{(t+1)} = \arg\min_{\boldsymbol{\gamma}} \left\{ \frac{1}{2m} \sum_{j=1}^{m} (\hat{\alpha}_j - \hat{\boldsymbol{\beta}}_j^\top \boldsymbol{\theta}^{(t+1)} - \gamma_j)^2 + \sum_{j=1}^{m} q_\tau(\gamma_i) \right\}. \tag{17}$$

We employ the alternating direction method of multipliers (ADMM) [1] to minimize (17). Specifically, we use the MCP as $q_{\boldsymbol{\lambda}_\gamma}$, which is expressed as:

$$q_{\tau,a}^{\text{MCP}}(x) = \tau \int_0^{|x|} \left(1 - \frac{|x|}{a\tau}\right)_+ dx \tag{18}$$

where $(x)+ = \max(x, 0)$ and $a > 1$ is an alternative tuning parameter that controls the concavity of $q_{\tau,a}^{\text{MCP}}(\cdot)$. When we denote $q_{\tau,a}^{\text{MCP}}(\boldsymbol{x})$ for a multivariate vector $\boldsymbol{x} = (x_1, \ldots, x_p)^\top$, it refers to

$$q_{\tau,a}^{\text{MCP}}(\boldsymbol{x}) = \sum_{j=1}^{p} q_{\tau,a}^{\text{MCP}}(x_j). \tag{19}$$

In this paper, we fix $a = 3$, as suggested by the original author, making $\tau$ the sole parameter in MCP. Second, the ADMM framework introduces an additional parameter, $\boldsymbol{\gamma}_1$, into (17), transforming it into

$$\boldsymbol{\gamma}^{(t+1)} = \arg\min_{\boldsymbol{\gamma} \in \mathbb{R}^m} \left\{ \frac{1}{2m} \|\hat{\boldsymbol{\alpha}} - \hat{\mathbf{B}} \boldsymbol{\theta}^{(t+1)} - \boldsymbol{\gamma}\|_2^2 + q_\tau(\boldsymbol{\gamma}_1) \right\}, \quad \text{subject to } \boldsymbol{\gamma} = \boldsymbol{\gamma}_1, \tag{20}$$

where $\hat{\boldsymbol{\alpha}} = (\alpha_1, \ldots, \alpha_m)^\top$ and $\hat{\mathbf{B}} = (\hat{\boldsymbol{\beta}}_1, \ldots, \hat{\boldsymbol{\beta}}_m)^\top$. Then ADMM uses the augmented Lagrangian method to handle it, which results in the following Q-function:

$$Q(\boldsymbol{\gamma}, \boldsymbol{\gamma}_1, \boldsymbol{u}) = \frac{1}{2m} \|\hat{\boldsymbol{\alpha}} - \hat{\mathbf{B}} \boldsymbol{\theta}^{(t+1)} - \boldsymbol{\gamma}\|_2^2 + q_\tau(\boldsymbol{\gamma}_1) + \boldsymbol{u}^\top (\boldsymbol{\gamma} - \boldsymbol{\gamma}_1) + \frac{\rho}{2} \|\boldsymbol{\gamma} - \boldsymbol{\gamma}_1\|_2^2, \tag{21}$$

where $\boldsymbol{u}$ represents the augmented Lagrangian multiplier and $\rho$ is an additional tuning parameter introduced in ADMM. The ADMM algorithm updates $\boldsymbol{\gamma}$, $\boldsymbol{\gamma}_1$, and $\boldsymbol{u}$ sequentially as follows:

$$\boldsymbol{\gamma}^{(t+1)} = \arg\min_{\boldsymbol{\gamma}} \; Q(\boldsymbol{\gamma}, \boldsymbol{\gamma}_1^{(t)}, \boldsymbol{u}^{(t)}), \tag{22}$$

$$\boldsymbol{\gamma}_1^{(t+1)} = \arg\min_{\boldsymbol{\gamma}_1} \; Q(\boldsymbol{\gamma}^{(t+1)}, \boldsymbol{\gamma}_1, \boldsymbol{u}^{(t)}), \tag{23}$$

$$\boldsymbol{u}^{(t+1)} = \arg\min_{\boldsymbol{u}} \; Q(\boldsymbol{\gamma}^{(t+1)}, \boldsymbol{\gamma}_1^{(t+1)}, \boldsymbol{u}). \tag{24}$$

Each update has a close-form solution. For (22), the score function is

$$\hat{\boldsymbol{\alpha}} - \widehat{\mathbf{B}}\boldsymbol{\theta}^{(t+1)} + \boldsymbol{\gamma}^{(t+1)} + \boldsymbol{u}^{(t)} + \rho(\boldsymbol{\gamma}^{(t+1)} - \boldsymbol{\gamma}_1^{(t)}) = 0, \tag{25}$$

and hence

$$\boldsymbol{\gamma}^{(t+1)} = \frac{1}{1+\rho}\left(\hat{\boldsymbol{\alpha}} - \widehat{\mathbf{B}}^{\text{PLS}}\boldsymbol{\theta}^{(t+1)} - \boldsymbol{u}^{(t)} + \rho\boldsymbol{\gamma}_1^{(t)}\right). \tag{26}$$

As for (23), the score function is

$$q'_\tau(\boldsymbol{\gamma}_1^{t+1}) + \boldsymbol{u}^{(t)} + \rho(\boldsymbol{\gamma}_1^{(t+1)} - \boldsymbol{\gamma}^{(t+1)}) = 0, \tag{27}$$

and hence

$$\boldsymbol{\gamma}_1^{t+1} = \text{mcp}_{\rho^{-1}\tau}(\boldsymbol{\gamma}^{(t+1)} + \rho^{-1}\boldsymbol{u}^{(t)}), \tag{28}$$

where

$$\text{mcp}_\tau(x) = \begin{cases} \frac{a}{a-1}\texttt{soft}_\tau(x), & \text{if } |x| \le \tau a, \\ x, & \text{if } |x| > \tau a, \end{cases} \tag{29}$$

where $\texttt{soft}_\tau(x) = \texttt{sign}(x)(|x| - \tau)_+$ is known as the soft-thresholding operator and $a = 3$. As for (24), the gradient update is applied:

$$\boldsymbol{u}^{(t+1)} = \boldsymbol{u}^{(t)} - \rho(\boldsymbol{\gamma}^{t+1} - \boldsymbol{\gamma}_1^{t+1}). \tag{30}$$

Note that $\hat{\boldsymbol{\gamma}}$ will not be strictly sparse. Therefore, we use $\text{sign}(\hat{\gamma}_1) \times \hat{\boldsymbol{\gamma}}$ as the final output once the entire minimization process converges.

The optimal choice of ADMM parameter $\rho$ is related to the spectral norm of Hessian matrix of the optimization; see, e.g., Theorem 1 of Ghadimi et al. [3]. In MRBEE-TL, the Hessian matrix of $\boldsymbol{\gamma}$ is just $\mathbf{I}$ whose eigenvalues are all 1. Therefore, it is reasonable to fix $\rho = 3$ and other values slightly larger than 1. In MRBEE-TL, the default choice of $\rho$ is 3 but we allow the user to tune it.

Note that in MRBEE-TL, we will indeed reweight the input effect sizes by:

$$\tilde{\alpha}_j \leftarrow \frac{\hat{\alpha}_j}{\text{se}(\hat{\alpha}_j - \alpha_j)}, \tag{31}$$

which makes

$$\text{E}(\tilde{\alpha}_j) = \frac{\sum_{s=1}\beta_{js}\theta_s + \gamma_j}{\text{se}(\hat{\alpha}_j - \alpha_j)}. \tag{32}$$

As a consequence, if an IV is invalid with $\gamma_j \neq 0$, its scale diverges at a rate of $1/\text{se}(\hat{\alpha}_j - \alpha_j)$, which is equal to $\sqrt{n_0}$. Hence, for selecting non-zero $\gamma_j$, the tuning parameter $\tau$ can be set as a sequence of 3,4,5,6,7 and others moderate large values. We do not suggest small $\tau$ as the background estimation error variance is

$$\text{var}(\tilde{\alpha}_j - \text{E}(\tilde{\alpha}_j)) = \frac{(\boldsymbol{\theta}^\top, -1)^\top \boldsymbol{\Sigma}_{xy}(\boldsymbol{\theta}^\top, -1)}{\text{se}(\hat{\alpha}_j - \alpha_j)} \tag{33}$$

which is usually around 1.

As for the correlation matrix of estimation error $\mathbf{\Sigma}_{xy}$, its estimation has been introduced by previous papers; see, e.g., Lorincz-Comi et al. [5]. Here we give a brief introduction. Specifically, two methods are commonly used to estimate the covariance matrix of estimation error $\mathbf{\Sigma}_{xy}$: LD score regression [2] and null effect estimate [11]. Here, we suggest estimating $\mathbf{\Sigma}_{xy}$ from statistically insignificant effect estimates because it is more straightforward and computationally efficient than LD score regression. Let $F_{i1}, \ldots, F_{iM}$ be $M$ independent genetic variants that are not associated with a trait with GWAS effect estimate $\hat{b}_{jk} = \sum_{i=1}^{n_k} F_{ij} X_{ik}/n_k$ and $\hat{b}_{j0} = \sum_{i=1}^{n_0} F_{ij} y_i/n_0$. Lorincz-Comi et al. [5] proved that $\hat{b}_{jk}$ and $\omega_{jk}$ has the same asymptotic distribution, allowing us to estimate $\mathbf{\Sigma}_{xy}$ by

$$\widehat{\mathbf{\Sigma}}_{xy} = \frac{1}{M} \sum_{j=1}^{M} \hat{\boldsymbol{b}}_j \hat{\boldsymbol{b}}_j^\top, \tag{34}$$

where $\hat{\boldsymbol{b}}_j = (\hat{b}_{j1}, \ldots, \hat{b}_{jp}, \hat{b}_{j0})^\top$. To ensure that the statistically insignificant variants used for calculating $\widehat{\mathbf{\Sigma}}_{xy}$ are approximately independent, we use a subsampling strategy. We start by randomly sampling a minor fraction (e.g., 10%) of variants from $M$ to estimate $\widehat{\mathbf{\Sigma}}_{xy}$, then repeat this process multiple times (e.g., 300), and finally use the mean of these estimates as $\widehat{\mathbf{\Sigma}}_{xy}$. Since LD decreases rapidly with increasing physical distance, each randomly sampled variants are nearly uncorrelated. Moreover, there are millions of common variants across the genome, the vast majority of which are not associated with a trait. Hence, sampling 10% or other minor fraction of the statistically insignificant variants can reliably yield $\widehat{\mathbf{\Sigma}}_{xy}$.

#### 3.5 Extended Bayesian information criterion

We used the EBIC to selecting the optimal parameters $L$ and $\tau$. The minimization of EBIC is

$$L^{\text{opt}}, \tau^{\text{opt}} = \arg\min_{L,\tau} \left\{ \log \hat{\sigma}^2(L,\tau) + \frac{(\log m + \text{ebic}_L \log p)||\hat{\boldsymbol{\delta}}(L)||_0 + (1 + \text{ebic}_\tau) \log m ||\hat{\boldsymbol{\gamma}}_1(\tau)||_0}{m} \right\}, \tag{35}$$

where

$$\hat{\sigma}^2(L,\tau) = \frac{\|\hat{\boldsymbol{\alpha}} - \hat{\mathbf{B}}\hat{\boldsymbol{\theta}} - \hat{\boldsymbol{\gamma}}\|_2^2}{m - ||\hat{\boldsymbol{\delta}}(L)||_0 - ||\hat{\boldsymbol{\gamma}}_1(\tau)||_0}, \tag{36}$$

$\text{ebic}_L$ and $\text{ebic}_\tau$ are two EBIC factors controlling additional penalities on the degrees of freedom of $\hat{\boldsymbol{\theta}}$ and $\hat{\boldsymbol{\gamma}}$. From a statistical perspective, as $p$ or $m$ increases substantially (i.e., diverges), it is appropriate for $\text{ebic}_L$ or $\text{ebic}_\tau$ to increase accordingly from zero. In this context, we recommend setting $\text{ebic}_L = 0$ and increasing $\text{ebic}_\tau$ to 1 or 2 when dealing with a large number of instruments. In addition, our default setting of the candidate set of $L$ is $\{1, 2, 3, 4, 5, 6, 7, 8\}$ and the default setting of candidate set of $\tau$ is $\{3, 3.5, 4, \ldots, 8\}$.

#### 3.6 Estimation of covariance matrix via stability selection

We apply the selection stability method [6] to estimate the covariance matrix and also empirical PIPs of $\hat{\boldsymbol{\theta}}$. Let $H$ be the number of sampling times and $\mathcal{I}^{(h)}$ be a random subsample of $1, \ldots, m$ of size $[0.5m]$ drawn without replacement. According to Meinshausen and Bühlmann [6], the sample size of $[0.5m]$ is chosen as it resembles most closely the bootstrap while allowing computationally efficient implementation. Next, we estimate $\hat{\boldsymbol{\theta}}^{[h]}$ and $\hat{\boldsymbol{\gamma}}^{[h]}$ by:

$$\begin{aligned}(\hat{\boldsymbol{\theta}}^{[h]}, \hat{\boldsymbol{\gamma}}^{[h]}) = \arg\min_{\boldsymbol{\theta},\boldsymbol{\gamma}} \Bigg\{ &\frac{1}{m} \sum_{j \in \mathcal{I}^{(h)}} (\hat{\alpha}_j - \hat{\boldsymbol{\beta}}_j^\top \boldsymbol{\theta} - \gamma_j)^2 - \frac{1}{2} \boldsymbol{\theta}^\top \mathbf{\Sigma}_{W_\beta W_\beta} \boldsymbol{\theta} + \boldsymbol{\sigma}_{W_\beta w_\alpha}^\top \boldsymbol{\theta} \\ &+ \sum_{j \in \mathcal{I}^{(h)}} q_{\tau^{\text{opt}}}(\gamma_i) + \sum_{k=1}^{p} \rho_{\lambda^{\text{opt}}}(\theta_k, 0, \hat{\theta}_{\text{source } k}) \Bigg\},\end{aligned} \tag{37}$$

where $\lambda^{\text{opt}}$ and $\tau^{\text{opt}}$ are the optimal ones determined by EBIC. We implemented such procedures for $H$ times and obtained a sequence of estimates $\hat{\boldsymbol{\theta}}^{[1]}, \ldots, \hat{\boldsymbol{\theta}}^{[H]}$. The covariance matrix and PIPs of this sequence were used as the covariance matrix and PIPs of $\hat{\boldsymbol{\theta}}_{\text{BEE-TL}}$.

##### 3.7 Supplementary Figures

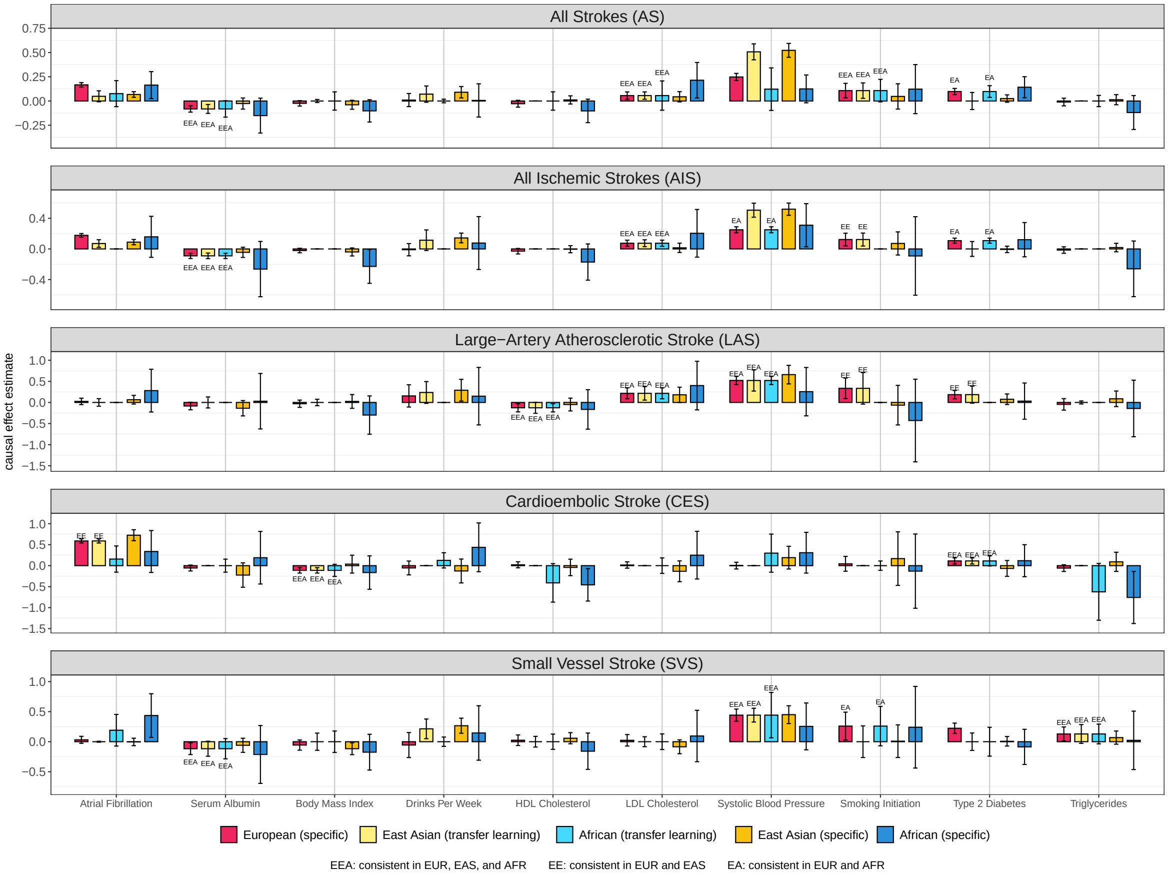

Figure S2: Causal effect estimates of selected exposures on five stroke outcomes across European, East Asian, and African ancestries. Each exposure-outcome pair is evaluated using five settings: European (specific), East Asian (transfer learning), African (transfer learning), East Asian (specific), and African (specific) (by using the original MRBEE). Bar heights represent point estimates of causal effects, with error bars indicating 95% confidence intervals. Annotations “EA", “EE", and “EEA" above bars denote consistent causal effect estimates between European and African, European and East Asian, and all three ancestries, respectively, identified by transfer learning. This figure shows the results using all IVs yielded by FEMA, without excluding any potential IVs with GGI or GEI evidence. The total number of IVs is 3194.

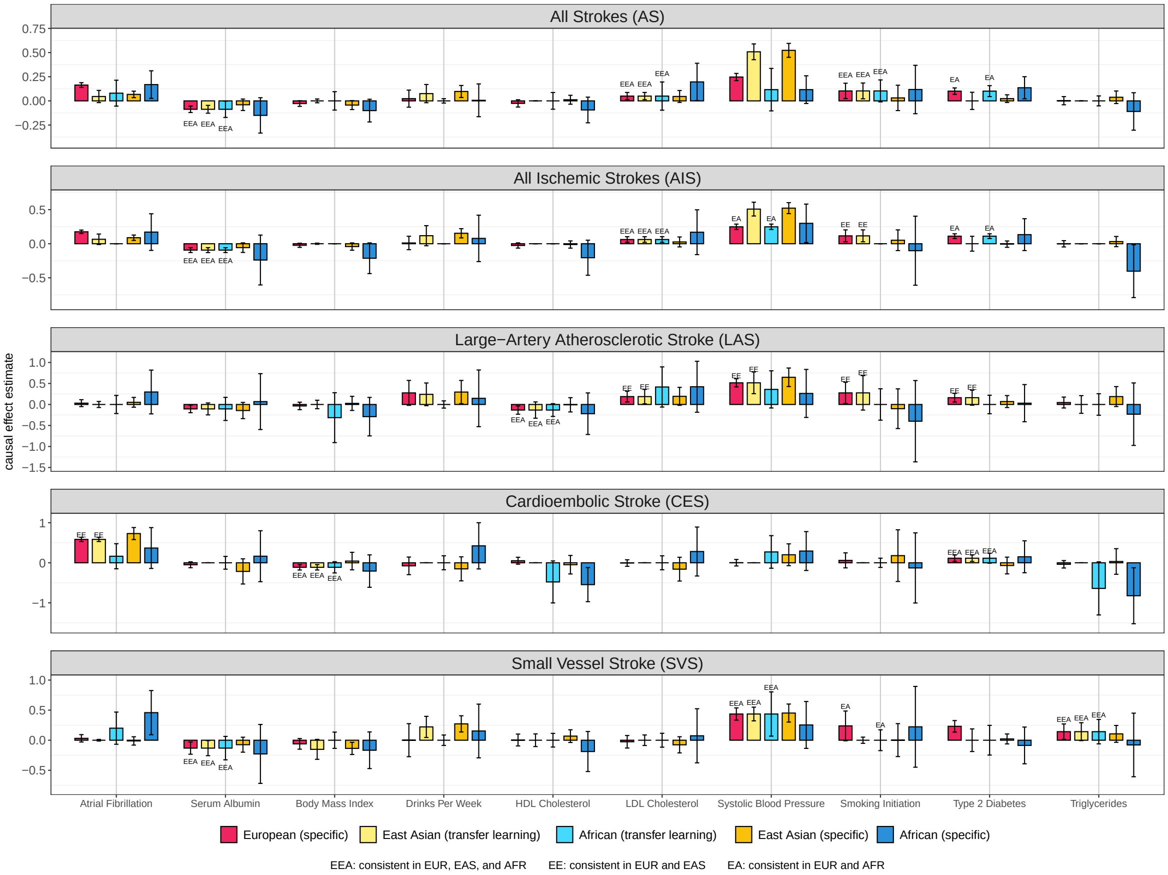

Figure S3: Causal effect estimates of selected exposures on five stroke outcomes across European, East Asian, and African ancestries. Each exposure-outcome pair is evaluated using five settings: European (specific), East Asian (transfer learning), African (transfer learning), East Asian (specific), and African (specific) (by using the original MRBEE). Bar heights represent point estimates of causal effects, with error bars indicating 95% confidence intervals. Annotations “EA", “EE", and “EEA" above bars denote consistent causal effect estimates between European and African, European and East Asian, and all three ancestries, respectively, identified by transfer learning. This figure shows the results where the IVs with potential GGI or GEI evidence were detected by MR GxE framework. The total number of IVs is 3150.
